# Inferring Dynamic Functional Connectivity from Field Potentials Using Graph Diffusion Autoregression

**DOI:** 10.1101/2024.02.26.582177

**Authors:** Felix Schwock, Julien Bloch, Karam Khateeb, Jasmine Zhou, Les Atlas, Azadeh Yazdan-Shahmorad

## Abstract

Estimating dynamic functional connectivity (dFC) is attracting increased attention, spurred by rapid advancements in multi-site neural recording technologies and efforts to better understand cognitive processes. Yet, most studies focus on static estimates of functional connectivity that cannot capture highly dynamic neural processes, while also ignoring information about the structural organization of the brain. To address these issues, we introduce a class of network-constrained linear autoregressive models that give rise to a highly dynamic functional connectivity signal on the edges of a predefined structural connectivity graph. Furthermore, we demonstrate that adding an additional diffusion constraint improves the model’s performance. We successfully validated the resulting graph diffusion autoregressive (GDAR) model on simulated neural activity and recordings from subdural and intracortical micro-electrode arrays placed in macaque sensorimotor cortex demonstrating its ability to describe rapid communication dynamics induced by optogenetic stimulation, changes in resting state dFC following stroke and electrical stimulation, and neural correlates of behavior during a reach task.

## Introduction

The coordinated interactions across different brain networks and subnetworks underlies cognitive processes^1–6^, and disruptions of these interactions are linked to a range of neurological disorders^7–10^. Despite this demonstrated importance, we still do not fully understand how brain networks perform computations through the coordinated signaling of connected neurons and neural populations during natural behavior, following a disease or injury, or as the result of rehabilitative intervention. The development of new electrophysiological recording technologies such as large-scale micro-electrode arrays provides unique opportunities for addressing this gap by measuring brain network activity simultaneously over multiple areas with high spatial and temporal resolution^11–16^.

A common signal extracted from subdural and intracortical micro-electrode arrays is the local field potential (LFP), which describes voltage fluctuations in the extracellular space of neuronal tissue. For these signals, the most common approach for estimating neural communication is through functional connectivity (FC) analysis^17,18^. In general, FC measures define neural communication as the undirected (symmetric) or directed (asymmetric) statistical dependence between different measurements that can be inferred from data using either model-free approaches or very general model classes such as vector autoregressive (VAR) models^19–22^. While these techniques are a popular choice for electrophysiology analysis, they predominantly yield static estimates of neural communication. Additionally, they rarely incorporate information about the structural network connectivity of the underlying brain region, particularly when analyzing recordings from high-resolution intracranial electrophysiology arrays.

To address these limitations, we propose a new approach for estimating dynamic functional connectivity (dFC) that 1) naturally incorporates the spatial layout of the recording array and the local connectivity structure of the cortex^23^ as a structural prior and 2) produces a highly dynamic and directed connectivity signal that can be used to study transient network events. Our approach is based on a class of network-constrained linear autoregressive models that can be fit to neural data. We consider two specific models of this class: 1) a standard vector autoregressive model whose parameters are constrained to a predefined structural connectivity graph called graph autoregressive (GAR) model and 2) a Laplacian autoregressive model, where network interactions are modeled via the graph Laplacian. Because the graph Laplacian is commonly used to model diffusion processes on networks^24^, we refer to the latter approach as the graph diffusion autoregressive (GDAR) model. Furthermore, we introduce new post processing techniques for analyzing the graph-spectral properties of network-constrained directed FC estimates that are based on the Hodge decomposition^25–28^.

To demonstrate the utility of our framework, we tested the GAR and GDAR model on five highly diverse datasets. First, using synthetic data from various networks of Wilson-Cowan oscillators we demonstrate that the high-resolution dFC signal estimated by the GDAR model better captures the simulated communication signal than GAR and standard VAR models. Furthermore, we show that GAR and GDAR models better generalize to unseen data than VAR models. Finally, using three micro-electrocorticography (*µ*ECoG) and one Utah array dataset we demonstrate that the GDAR model can be used to uncover transient communication dynamics evoked *during* cortical optogenetic stimulation, characterize changes in resting state FC *after* stroke and electrical stimulation, and uncover network-level dFC correlates of a monkey’s reach behavior that are dependent on the spatial frequency. We show that the GDAR model outperforms alternative approaches and provides insights that cannot be obtained by other models.

## Results

### Graph autoregressive (GAR) and graph diffusion autoregressive (GDAR) model

GAR and GDAR models are linear vector autoregressive (VAR) models, whose parameters are constrained to the connections of a predefined structural connectivity graph. An overview of both models is shown in Fig. 1 and a more detailed mathematical description can be found in Methods. First, the electrode layout of the recording array is used to construct a sparse and locally connected graph, with each electrode representing a node and with edges connecting nearby nodes Fig. 1a left). This graph serves as a structural prior that incorporates information about the local connectivity of the cortex into the model^23^. The models then transform the neural activity into directed FC or flow signals defined on the graph edges (Fig. 1a right). These signals, which we will refer to as *GAR flow* and *GDAR flow*, respectively, describe the predictive information flow between the nodes. Unlike classical FC analysis, which aggregates information over multiple time points, thereby estimating an average information flow, the GAR and GDAR models transform the neural activity at each time point into a flow signal without losing temporal resolution. Therefore, they can be used to study highly transient communication events in the brain.

**Fig. 1.**
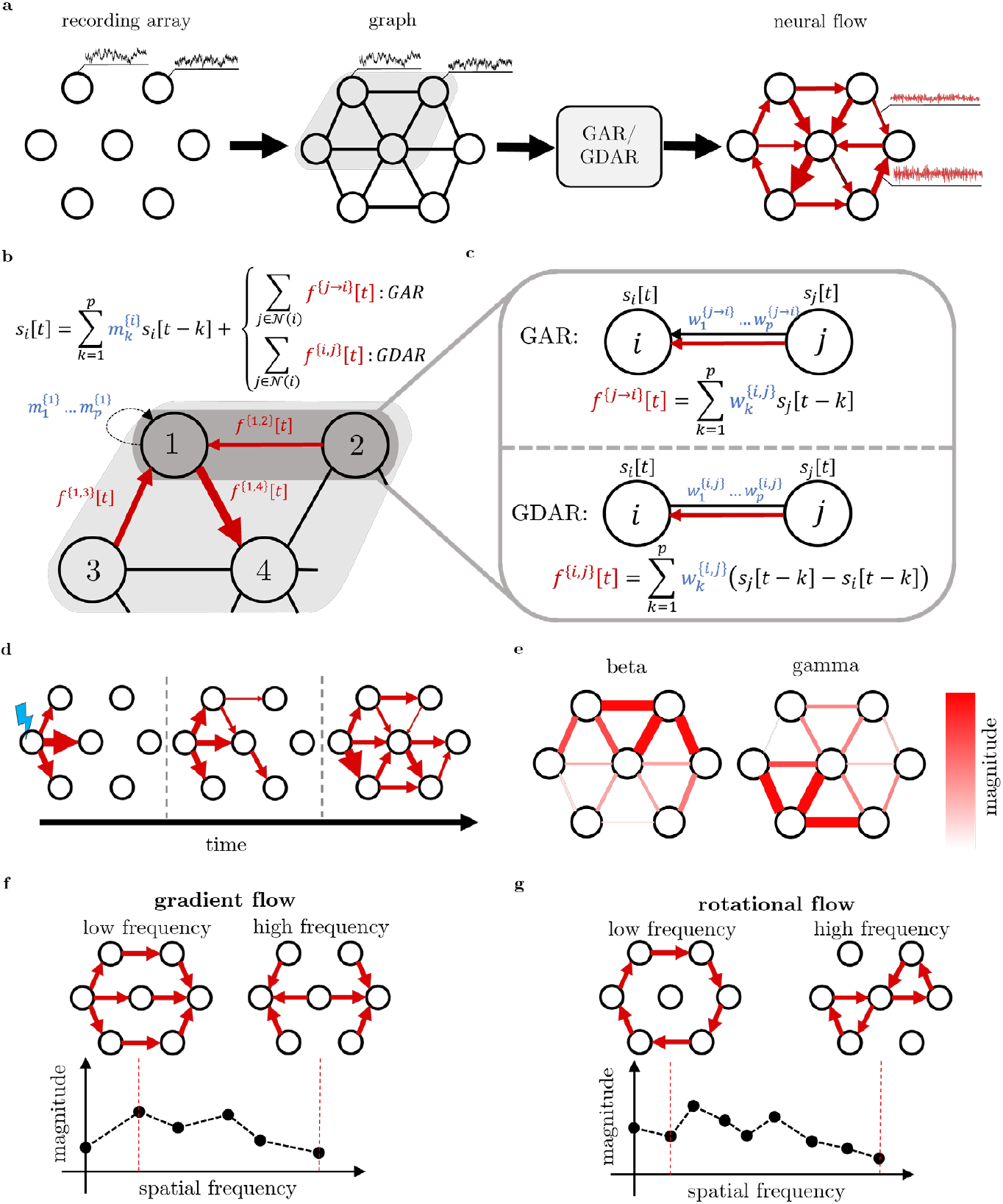
Overview of the graph autoregressive (GAR) and graph diffusion autoregressive (GDAR) model. (a) The recording array is used to form a sparse, locally connected graph, where each electrode represents a node, and edges connect neighboring nodes. GAR/GDAR models then transform the neural activity observed at the nodes into a directed flow signal defined on the graph edges, representing the real-time signaling between nodes. (b) The models incorporate an order p autoregressive system, where at time t each node’s neural activity is modeled using a combination of its past p samples and the flow from all adjacent nodes. (c) The GAR flow f^{j→i}^[t]from node j to iat time t is given by a weighted sum of the previous pp samples from node jj. The GDAR flow f^{i,j}^[t], is calculated as the weighted sum of the previous pp activity gradients between iand j. In analogy to current source density analysis, the edge parameters of the GDAR flow 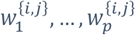 can be interpreted as conductivities for local field potential measurements, such that conductivity times potential gradient yields a current flow. The model parameters are assumed to be static within a particular time window and can be estimated using linear regression (see Methods). (d) The GAR/GDAR flow can be used to study transient communication events, for example due to cortical stimulation. (e) For resting state recordings, power spectral density estimates of the flow signals can be used to study frequency specific communication patterns. (f), (g) Akin to classical Fourier analysis for time series, the GDAR flow signal can also be decomposed into gradient (non-circular) and curl (circular) flow modes of different spatial frequency to study the smoothness and spatial composition of the flow signal across the network.

GAR and GDAR flow are obtained by deconstructing the prediction of a *p*^th^ order vector autoregressive model at time *t* and node *i*into its contribution from the past of node *i*itself, and the information provided by the neighbors of node *i*(Fig. 1b and c). Formally, let *si*[*t*]be the neural activity at node *i* and time *t*, then

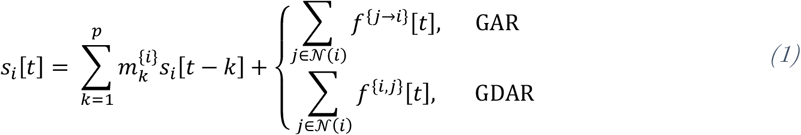

where 𝒩 (*i*) is the set of neighbors of node *i*. GAR flow *f*^{*j*→*i*}^[*t*]and GDAR flow *f*^{*i,j*}^[*t*]are given by

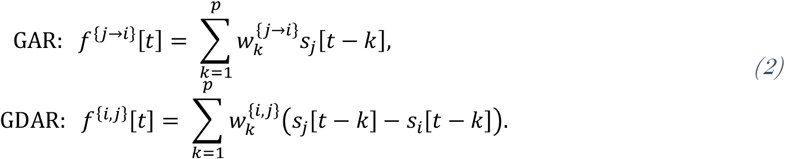

The main difference between the GAR and GDAR model is that the predictive information flow for the GDAR model is driven by gradients of activity, whereas for the GAR model it is driven by the activity itself. The GDAR flow can therefore be considered analogous to classical heat diffusion, where heat gradients drive changes in temperature. The idea of modeling neural interactions via a diffusive process has also been proposed in other neuroscience contexts^29,30^.

The parameters of the models 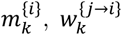, and 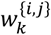 can be estimated from neural recordings using linear regression (see Methods) and are assumed to be static within a predefined time window. It is worth noting that the GDAR model only has one direction agnostic parameter 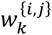 per edge and lag, whereas the GAR model uses direction specific weights 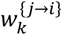 and 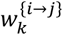 that are generally not equal. Since the activity gradient from node *i* to *j* is the negative of the activity gradient from *j* to *i*, a positive flow from node *j* to *i*(which is equivalent to a negative flow from *i* to *j*) causes an increase in activity at node *i*and an equal amount of decrease at node *j*. Therefore, the GDAR model produces a consistent flow across the network. On the other hand, the GAR model does not have this property as the flow from *j* to *i*does not affect the predicted activity at node *j*. We note that both GAR and GDAR flow are directed dFC estimates. However, while the GAR flow is bidirectional at each time point, where the flow in each direction can be positive or negative, the GDAR flow is unidirectional and can be interpreted as the *net flow* between two nodes. However, the direction of the GDAR flow across each edge can, and generally does, change over time.

For a model order *p* = 1 and constant weights *w*^{*i,j*}^, the GDAR model is equivalent to computing current source densities (CSDs), which is a popular technique for analyzing field potential recordings obtained from technologies such as ECoG or electroencephalography (EEG)^31,32^. Therefore, the GDAR model can also be considered a combination of CSD analysis and VAR model. For field potential recordings the model parameters 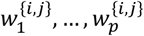 can be interpreted as conductivities such that voltage gradients multiplied by conductivities yields current flows. Summing the current flows at each node is analogous to computing the current sources and sinks in CSD analysis.

The high temporal resolution of the GAR and GDAR flow signal is ideal for studying transient signaling events. For example, the propagation of neural activity due to cortical stimulation can be tracked by analyzing the spatiotemporal evolution of consecutive time steps of *f*^{*i,j*}^[*t*]or *f*^{*j*→*i*}^[*t*](Fig. 1d). Alternatively, the models can be applied to resting state recordings by computing power spectra of the time varying GAR/GDAR flow resulting in a similar frequency decomposition that is typical for standard VAR based FC measures (Fig. 1e). Furthermore, modeling FC on top of a graph allows us to use recently developed theory from signal processing over simplicial complexes^25–27^ to decompose the flow signals into gradient (non-circular) and curl (circular) modes of different spatial frequencies (see Methods, Fig. 1f and g, and Extended Fig. 1 for examples). The resulting gradient and curl flow spectra can be used to quantify the degree of smoothness or coordination of the neural signaling (e.g., flow spectra with stronger low-frequency components are considered to represent more coordinated signaling). Since the decomposition requires unidirectional flow signals, this analysis tool is better suited for the GDAR model; however, it can also be adopted for the GAR model by converting the bidirectional GAR flow into a unidirectional flow (e.g., by computing *f*^{*j*→*i*}^[*t*]− *f*^{*i*→*j*}^[*t*]).

### The GDAR model outperforms VAR and GAR models in inferring communication dynamics in a network of coupled Wilson-Cowan oscillators

To assess the accuracy of the GAR/GDAR flow, we use simulated data generated by 10 randomly connected 16-node networks of coupled Wilson-Cowan oscillators (Fig. 2 and Extended Fig. 2a). The networks are used to generate a ground truth neural flow signal, as well as simulated neural activity, which is used to fit GAR, GDAR, and VAR models of varying model orders (see Methods). All models are used to transform the simulated neural activity into a neural flow signal, which is compared to the ground truth neural flow using various metrics (Fig. 2a). Furthermore, we have estimated traditional FC metrics via coherence or metrics from the Granger causality (GC) family. GC can be computed from the spectral representation of VAR model parameters leading to two metrics that are popular in neuroscience: partial directed coherence (PDC)^22^ and directed transfer function (DTF)^21^. For all VAR-based metrics (VAR flow, PDC, DTF), which assume fully connected model matrices, the resulting FC estimate is only compared over edges that are present in the structural connectivity networks.

**Fig. 2.**
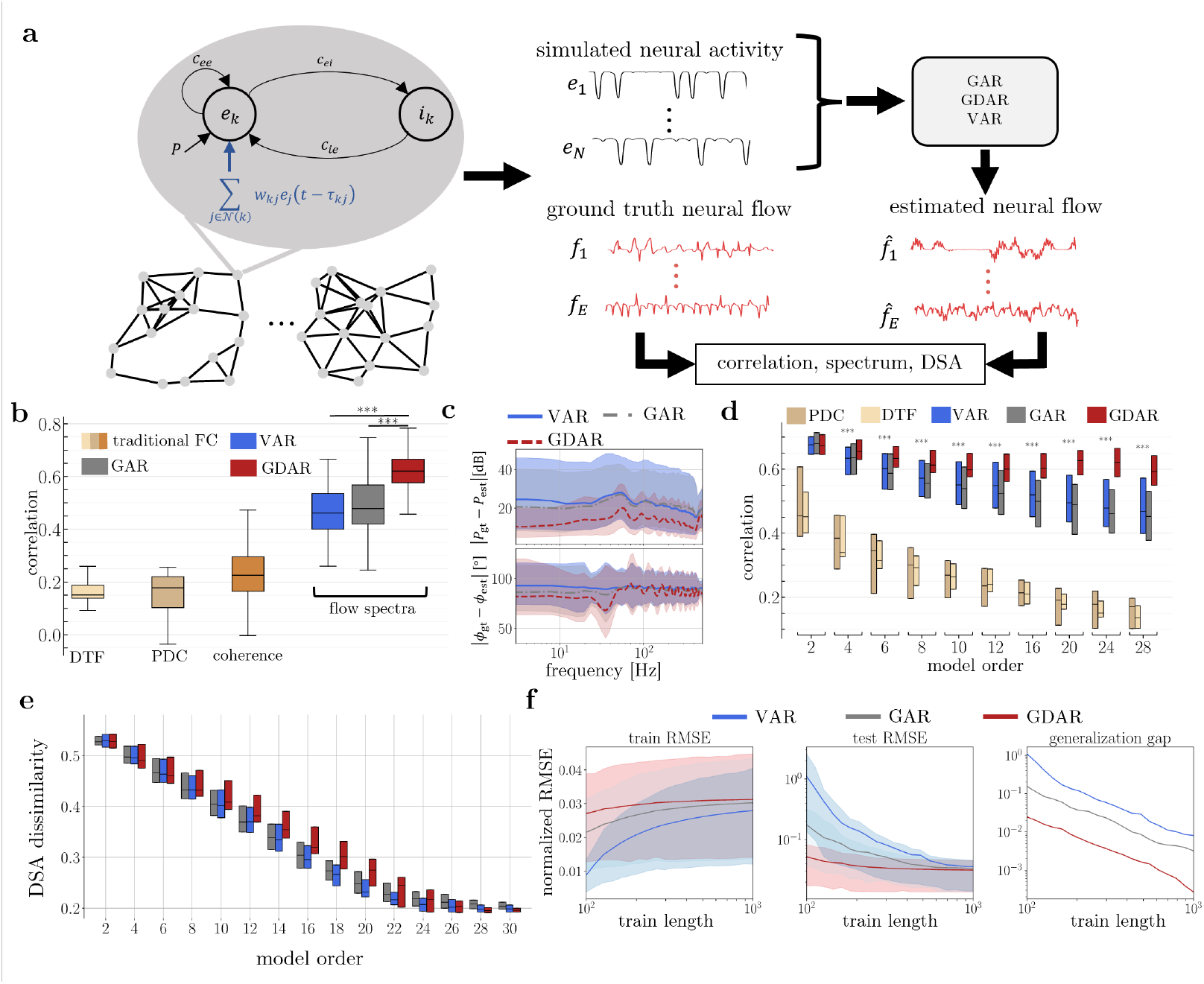
Evaluation of the GAR and GDAR model’s accuracy in capturing neural communication dynamics using networks of Wilson Cowan oscillators. (a) 10 randomly connected 16-node toy networks were used to simulate neural activity at each network node as well as compute a ground truth flow across each edge (see Methods). The estimated neural activity is transformed into the estimated neural flow signal using VAR, GAR, and GDAR models of different model orders, as well as the CSD approach. Ground truth and estimated neural flow are then compared using various metrics. (b) Distribution of Pearson correlation coefficients (medians, upper and lower quartiles, interquartile range) between ground truth flow spectrum and coherence, partial directed coherence (PDC), directed transfer function (DTF), and estimated flow spectrum pooled over all graph edges from 100 independent simulation trials (10 per network). The GDAR flow spectrum significantly outperforms all other metrics. (c) Magnitude and phase difference between the spectrum of ground truth and estimated flow (median, upper and lower quartile). The GDAR model shows consistently lower magnitude errors for all frequencies and phase errors below 50 Hz. (d) Same as (b) but for different model orders. The GDAR model significantly outperforms all other metrics for p ≥ 4. (e) Dissimilarity scores between estimated and ground truth flow signals obtained via dynamical similarity analysis (DSA) to assess the accuracy of the estimated flow dynamics. Low dissimilarity scores for high model orders (p ≥ 24) suggest an accurate estimation of the flow dynamics by all three models. (f) Generalization performance of 10^th^ order VAR, GAR, and GDAR models. All models were fit to the first NN samples (train length) of neural activity simulated by a 7-node network of Wilson-Cowan oscillators (see Extended Fig. 2e) independently for each of the 100 simulation trials. The model coefficients are then used to perform a one-step ahead prediction on the training data as well as the remaining samples in the trial (test data). The left and middle panel show the mean, 10^th^, and 90^th^ percentile of the root-mean-square prediction error (RMSEs). The generalization gap (right panel) is defined as the difference between mean test and train RMSE. The GDAR model generalizes significantly better to unseen data than the VAR and GAR model. All statistical tests use Wilcoxon rank-sum tests at a significance level p ≤ 0.05. (d) Significance markers compare GDAR with GAR model. (b)-(e) Exact p-values can be found in Supplementary Table 1 and Supplementary Table 2.

First, we assessed the accuracy of the estimate flow signal using spectral analysis for a single model order (*p* = 24). In Fig. 2b, we compared the spectral magnitudes of coherence, PDC, DTF, and estimated flow with the ground truth flow spectral magnitude using Peason correlation coefficients. We found that the GDAR flow yields significantly higher correlations than any other metric (Fig. 2d; Wilcoxon rank-sum test, *p* < 0.05). Since GDAR and GAR are based on autoregressive models, we can compute equivalent PDC and DTF metrics from their model parameters (see Methods). While GDAR-PDC/DTF outperform GAR- and traditional PDC/DTF they do not outperform any of the flow metrics (Extended Fig. 2d). Furthermore, we found that the GDAR model exhibits consistently lower magnitude errors than the VAR and GAR models for all frequencies and lower phase errors for frequencies below 50 Hz (Fig. 2c). These findings also generalize to other model orders. Specifically, for all *p* ≥ 4, correlations with the ground truth magnitude spectra were highest for the GDAR model (Fig. 2d) suggesting that it captures neural interactions more accurately than the alternative metrics. We found that the same observation also holds when computing correlation between estimated and ground truth flows in the time domain (Extended Fig. 2f).

A notable observation is that the spectral magnitude of the ground truth flow is well approximated by lower order VAR, GAR, and GDAR models. Specifically, the median correlation between estimated and ground truth spectral magnitude for VAR and GAR models decreases with increasing model order. In contrast, the correlation for the GDAR model reaches another local maximum at higher orders, where it significantly outperforms the other two models (Fig. 2d). Despite this second maximum, the highest median correlations for the GDAR model still occur at low model orders. At the same time, because low-order models rely on fewer past time steps for predicting future activity, they may have limited capacity to capture more complex spatiotemporal dynamics that are not fully reflected in spectral magnitude alone. To test this, we used a recently developed tool from dynamical systems theory, called dynamical similarity analysis (DSA)^33^, which uses dynamical mode decomposition and shape analysis to provide a dissimilarity score between two (high-dimensional) time series. We found that with increasing model order the accuracy in capturing the spatiotemporal dynamics improves for all models (decreased DSA dissimilarity score) before plateauing at an order around *pp* = 24 (Fig. 2e). Hence, to accurately capture the complex dynamic properties of the flow signal, higher order models are needed. For these higher orders, the GDAR model perform significantly better than VAR and GAR models at approximating the ground truth flow (Fig. 2b-d; Extended Fig. 2f).

Finally, we tested the models’ ability to generalize to unseen data by evaluating their one-step ahead prediction performance on data that were not included during model fitting. Using simulated neural activity from a 7-node network of Wilson-Cowan oscillators shown in Extended Fig. 2g (100 independent trials with 5s of simulated neural activity per trial), VAR, GAR and GDAR models were trained on the initial *N* samples of each trial and then tested on the remaining samples (see Methods). An advantage of the GDAR model, owing to its fewer parameters compared to both VAR and GAR models, is its reduced need for extensive training data to accurately estimate model parameters. This results in flatter root-mean-square error (RMSE) curves for both train and test set (Fig. 2f left and middle) as well as the lowest generalization gap (difference between test and train RMSE) among all models (Fig. 2f, right). Notably, the generalization gap for the GDAR model is nearly an order of magnitude smaller than that for the GAR model despite having only 37% fewer parameters. Similarly, the GDAR model provides near perfect generalization on a range of electrophysiology datasets and significantly outperforms GAR and VAR models especially when only limited data is available for model fitting (Extended Fig. 2h).

### Application of GDAR model to electrophysiological recordings

Due to better performance of the GDAR over the GAR model on simulated data, we will focus on the GDAR model for the remainder of the paper. To show the versatility of the GDAR model for analyzing network-level dFC evoked by cortical stimulation, during behavior, and at rest, we have applied the model to electrophysiological recordings from four separate experiments that either use µECoG arrays (3 datasets) or a Utah array (1 dataset) to record LFPs from the sensorimotor cortex of macaque monkeys. For all datasets, the layout of the recording array was first used to construct a locally connected and sparse graph—with nodes corresponding to the recording channels— mimicking the local connectivity structure of the cortex^23^. Next, GDAR models of different orders were fit to the recorded LFPs. The resulting model coefficients were then used to transform the LFPs into GDAR flow signals, which were post-processed depending on the experimental paradigm.

#### The GDAR model uncovers communication dynamics evoked by cortical optogenetic stimulation

First, we show that the GDAR model can uncover fast, stimulation induced communication dynamics that match the experimental protocol. To do so we used three sessions from an optogenetic stimulation experiment performed in macaques, where two lasers repeatedly stimulated different locations of the primary motor (M1) and somatosensory (S1) cortex expressing the opsin C1V1, and fit a 5^th^ order GDAR model to the LFPs recorded by a 96-channel µECoG array during stimulation (see Fig. 3a and b and Methods)^11,34,35^. GDAR flow signals averaged over all stimulation trials in the milliseconds before and after stimulation for Session 1 are shown in Fig. 3c and Supplementary Video 1. When the network is at rest, flow levels across the network are small. Activation by the first laser located in M1 causes the GDAR flow to immediately increase near the stimulation location before spreading further into the network and reaching S1. After the second laser was activated, the flow increases near the second stimulation location and spreads into most parts of the network within the next few milliseconds.

**Fig. 3.**
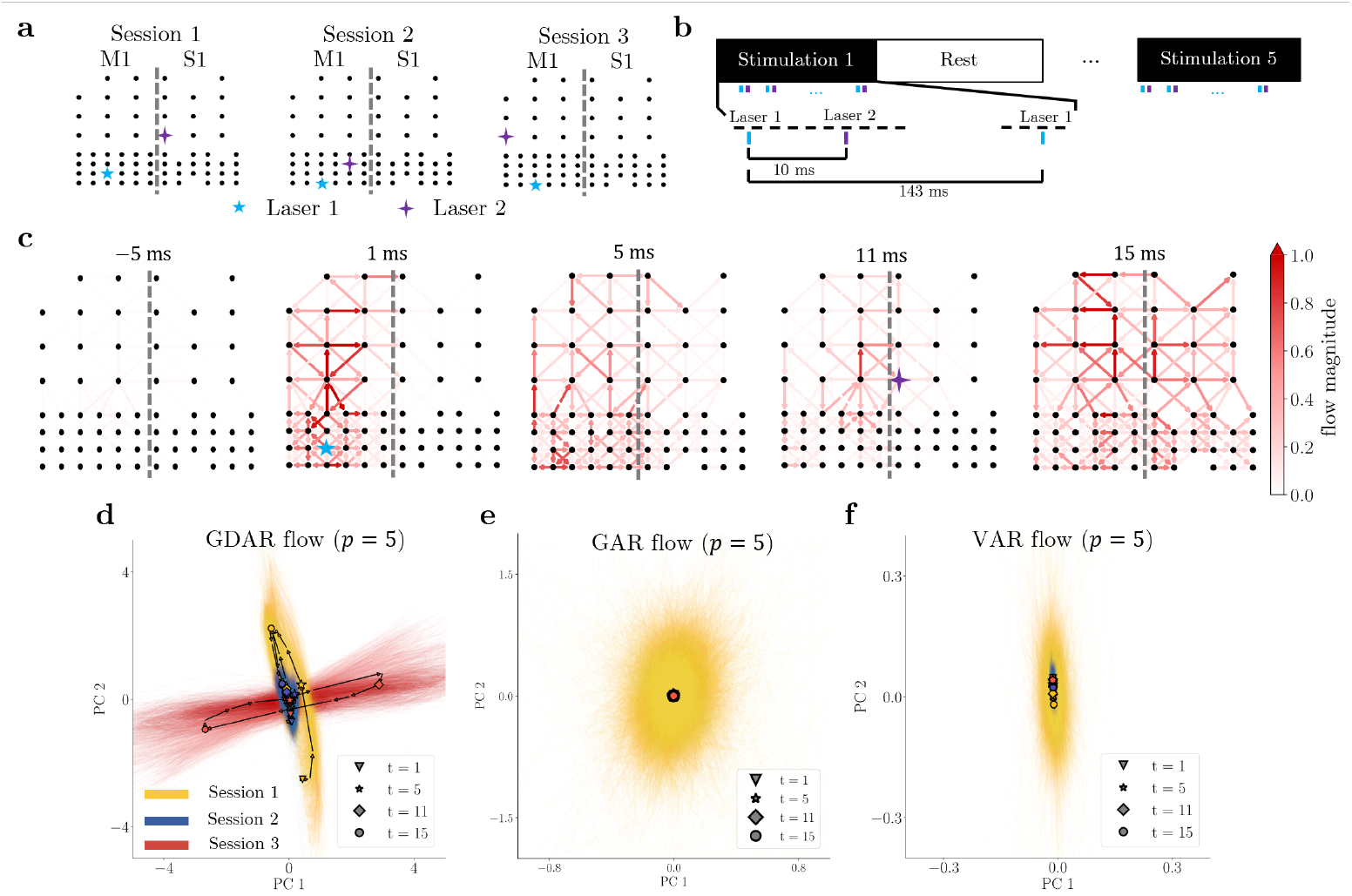
GDAR model applied to optogenetic stimulation experiment to study transient communication events. (a) LFPs from the primary motor (M1) and somatosensory (S1) cortex of a macaque monkey were recorded using a 96-channel µECoG array, while repeated paired stimulation was performed using two lasers. The relative positions of the electrodes after rejecting bad channels and the locations of the two lasers are shown for the three sessions that were analyzed in this work. The location of the sulcus between M1 and S1 is approximated by the thick gray line. The electrode array was not moved between the sessions. (b) Illustration of the stimulation protocol, where each laser stimulates alternatingly for 5 ms, with a 10 ms delay between stimulation by Laser 1 and 2. This paired stimulation is repeated every 143 ms. Each stimulation block lasts approximately 7 min and is intermitted by 20 s long resting blogs during which no stimulation is performed. (c) The GDAR flow for Session 1 averaged over all stimulation blocks and trials is shown for different time steps before (first plot) and after (remaining four plots) onset of stimulation from the first laser. The graphs suggest complex spatiotemporal signaling patterns evoked by cortical stimulation. (d) Flow snapshots from the first 25 ms after onset of the first laser stimulation for all trials, blocks, and sessions were stacked into a single matrix and the flow snapshots were projected onto its first two principal components (PCs). The PC reduced GDAR flow trajectories for different sessions are indicated by different colors. Average trajectories are shown as black solid lines with markers indicating different times point after the onset of stimulation by the first laser. Thin colored lines show trajectories by individual paired pulse trials. The GDAR flow trajectories are very consistent within and distinct between sessions, demonstrating that transient flow dynamics depend on the stimulation parameters. (e), (f) PC reduced flow trajectories as in (d) but using 5^th^ order GAR and VAR models. In contrast to the GDAR flow, GAR and VAR flow do not exhibit significant time and session dependent dynamics, thus, highlighting the utility of the GDAR model in uncovering stimulation induced transient communication dynamics.

It is apparent from Fig. 3c that the GDAR flow exhibits complex spatiotemporal dynamics within milliseconds after stimulation. To test how these dynamics depend on the stimulation pattern, we project the high-dimensional flow signal from three sessions, which only differ in their stimulation location (see Fig. 3a), onto their first two principal components (PCs) and compare the flow dynamics in this lower dimensional subspace (see Methods). We found that these low dimensional GDAR flow dynamics are very consistent within each session and strongly differ between sessions (Fig. 3d). Furthermore, the flow dynamics show some remarkable similarities with the stimulation patterns. For Sessions 1 and 2, where the flow trajectories largely align in the PC space, the stimulation patterns are similar in that the second stimulation occurs to the top right of the first stimulation. On the other hand, for Session 3, which results in flow trajectories orthogonal to Session 1 and 2, the second stimulation occurs to the top left of the first stimulation. Furthermore, the magnitude of the PC reduced GDAR flow dynamics is noticeably smaller for Session 2 compared to Sessions 1 and 3. This might be a result of the spatial separation between Laser 1 and 2, which is smallest for Session 2. Our findings extend previous work showing that LFP power in monkeys and humans distinctly depend on stimulation parameters such as amplitude and frequency^36,37^.

We also tested whether the GAR or VAR flow, or node signal like raw LFPs or its second spatial derivative, which resembles traditional current source density, can uncover dynamics that depend on the stimulation pattern but found that this is not the case (Fig. 3e-g, and Extended Fig. 3a-c). Perhaps it is not surprising that the raw LFPs or simple, model-free transformations thereof (e.g., second spatial derivative of raw LFP) fail to describe stimulation dependent dynamics using PC analysis as these signals may be dominated by noise and non-stimulation specific variation. In contrast, autoregressive models may effectively filter out some of these non-stimulation noise sources. Our results suggest that the GDAR model is more effective at uncovering such transient stimulation-dependent communication dynamics compared to alternative autoregression based models. We also note that the dependence on the stimulation location can be observed when plotting a low-dimensional representation of the model parameters itself, where the GDAR model shows a stronger separation between sessions than the VAR model (Extended Fig. 3g and h). Finally, the model can also be adapted to model longer signal propagation paths between specific nodes in the network as it may be the case for connections across the sulcus between M1 and S1 (see Extended Fig. 3f, Supplementary Video 2, and Supplementary Note).

#### The GDAR model can characterize network and graph-spectral changes in resting state FC that are complementary to previous FC techniques

One of the most common applications of FC analysis is comparing functional network connectivity across resting state recordings—for example, to assess changes related to mental states, disease, injury, or rehabilitation. While the GDAR model was primarily designed to resolve fine temporal communication dynamics, here we demonstrate using LFP data from two distinct experiments that the GDAR model is also well suited for analyzing frequency-specific resting state FC networks and that it can provide complementary insights to existing approaches. The first dataset contains cortical surface field potentials recorded using two 32-channel µECoG arrays before and after the induction of an ischemic lesion as well as following cortical stimulation in the acute phase after stroke (Fig. 4a and b). The second dataset contains intercortical recordings by a 96-channel microelectrode array (Utah array) before and after repeated electrical stimulation (Fig. 5). All experiments were performed in the non-human primate sensorimotor cortex. All model-based techniques use a model order of 10 and connectivity/flow changes are tested for statistical significance using a two-sample Kolmogorov-Smirnov test (see Methods).

**Fig. 4.**
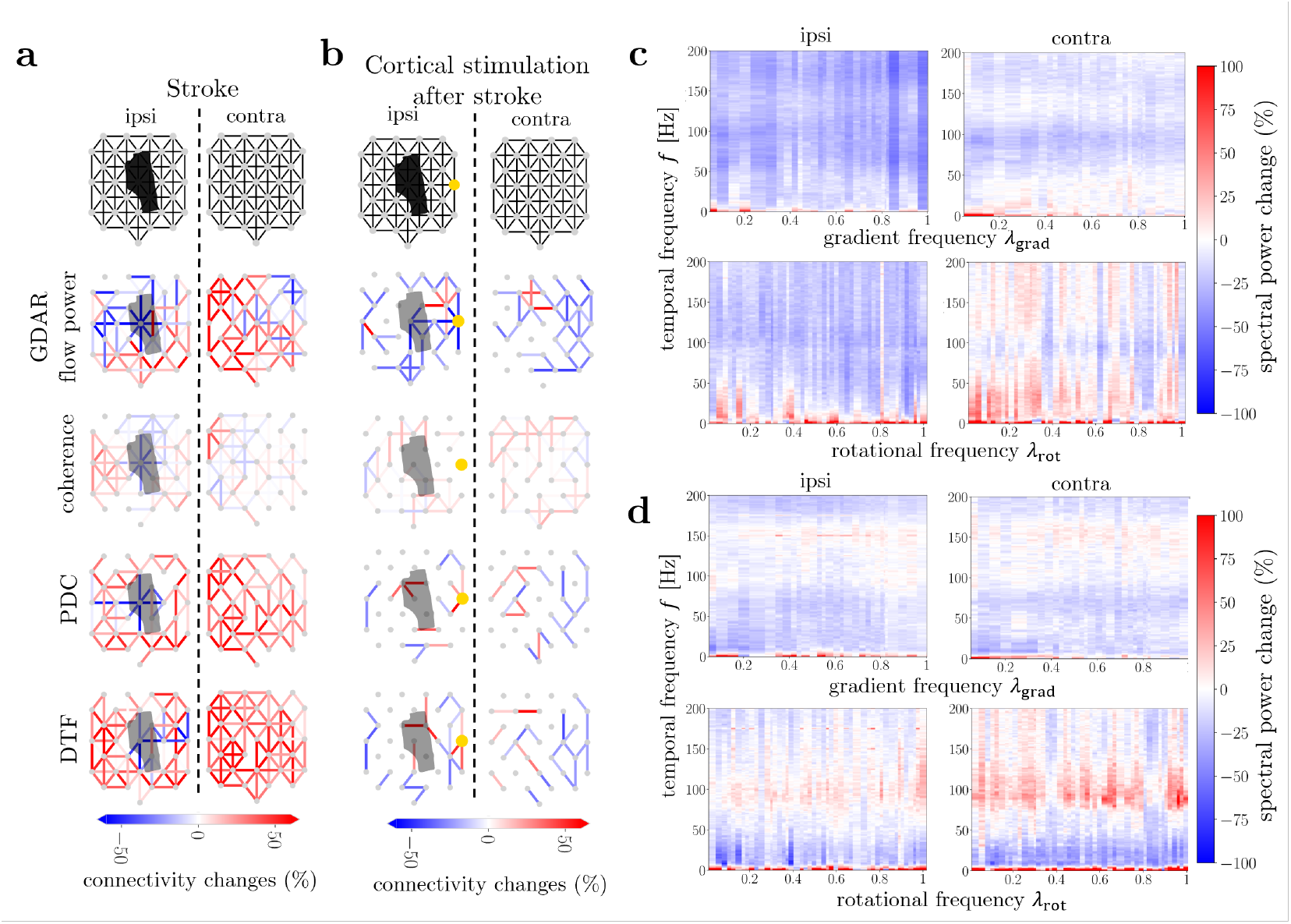
Application of GDAR model for resting-state FC analysis using LFP data recorded from macaque sensorimotor cortex during an ischemic lesion and cortical stimulation experiment. (a) Changes in gamma (30-70 Hz) GDAR flow power, coherence, PDC, and DTF due to stroke in ipsilesional (left) and contralesional (right) hemisphere. The first column shows the relative locations of the recording electrodes and corresponding structural connectivity graphs. The black patch indicates the extent of the cortical lesion for the stroke experiment. (b) Same as (a) but now showing changes following electrical stimulation in the acute phase after stroke. Yellow markers show the location of electrical stimulation. Only edges with significant changes are shown (p<0.01, two-sample Kolmogorov-Smirnov test). (c), (d) Changes in spatiotemporal spectral power due to (c) stroke and (d) cortical stimulation after stroke. Spatiotemporal spectra are calculated by first computing the time varying gradient and curl spectrum for each time step of the GDAR flow followed by estimating the time power spectral density of the gradient and curl spectral coefficient time series using Welch’s method. In general, the curl flow exhibits an increase in spectral power over larger spatial and temporal frequency ranges for both hemispheres suggesting an increase in feedback communication. For all model-based techniques a model order of p = 10 was used.

**Fig. 5.**
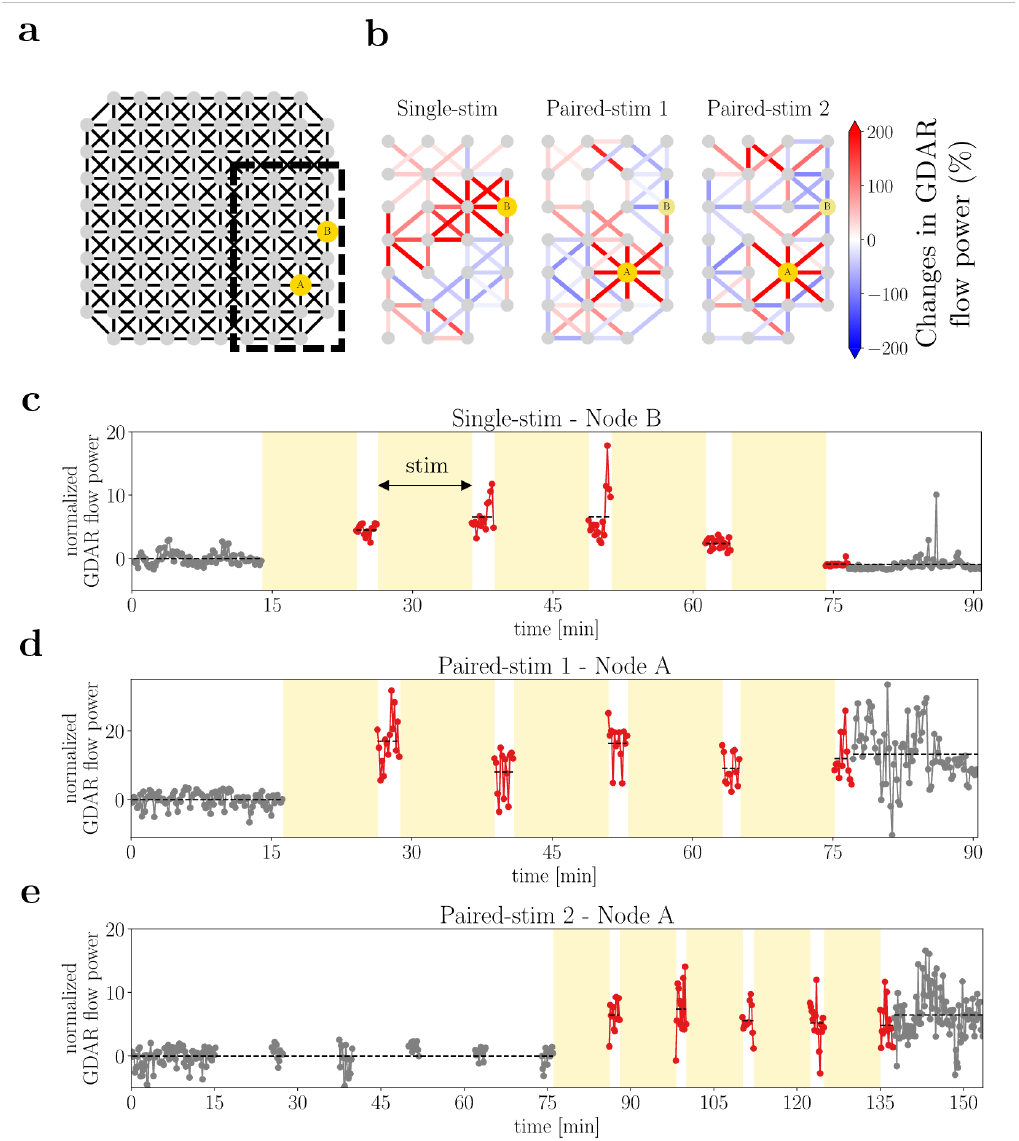
Application of GDAR model to LFP data recorded from a macaque monkey using a Utah array during a paired electrical stimulation experiment. (a) The locations of the electrodes were used to construct a locally connected sparse graph as input to the GDAR model. Electrode A and B are used for single-site and paired stimulation. For paired stimulation, electrode A stimulates before electrode B. (b) Changes in gamma (30-70 Hz) GDAR flow power due to stimulation are shown for three separate sessions. For the single-stim session, only electrode B stimulates. For the paired-stim sessions, electrode A and B stimulate repeatedly and alternatingly for a total of 50 minutes divided into five 10min blocks (see Methods for more details on the stimulation protocol). An increase in gamma GDAR flow power near the first stimulation location can be observed for all sessions. (c)-(e) Temporal evolution of the normalized gamma GDAR flow power averaged over all edges adjacent to the first stimulation site for all three sessions. The 2-minute blocks immediately after stimulation are highlighted in red. The gamma GDAR flow power has been corrected for linear changes in LFP power at the stimulation site (see Methods). For the paired stimulation sessions, the GDAR flow power remains elevated even after the stimulation period ends.

For the ischemic stroke experiment we qualitatively compare changes in gamma (30-70 Hz) GDAR flow power across the ipsi- and contralesional hemispheres with changes in coherence, PDC, and DTF. All methods showed decreased gamma connectivity for edges within the lesioned area consistent with the occurrence of neuronal death (Fig. 4a). In perilesional and contralesional areas, all methods show increased FC consistent with previous findings^38,39^; however, the location and extent of decreased FC vary greatly between techniques. Electrical stimulation in the acute phase after stroke (Fig. 4b) caused a decrease in GDAR flow power across most of the ipsi- and contralesional network, particularly for edges connected to the stimulation site. This is consistent with previous findings from our lab demonstrating that electrical stimulation reduces post-stroke neural and c-Fos activity^40,41^. In contrast, PDC and DTF show more mixed connectivity changes, while coherence shows increased connectivity across both hemispheres. Taken together, these results reveal broad qualitative agreement between GDAR flow, PDC, and DTF in characterizing FC changes due to stroke and cortical stimulation. Nonetheless, the GDAR model also captures changes not detected by the other methods, suggesting it can provide complementary insights into resting-state FC analysis.

In addition, GDAR inherently models FC on top of a structural graph that allows for performing a graph spectral analysis as illustrated in Fig. 1f and g and Extended Fig. 1. Specifically, at each time point, we can compute the gradient and curl flow spectrum *F_grad_* and *F_curl_* from the GDAR flow signal *f*, which describe the magnitude of non-circular (gradient) and circular (curl) flow patterns, where each pattern exhibits different degrees of spatial smoothness. Similar to the GDAR flow, the gradient and curl flow spectra are time varying resulting in a time series for each spectral coefficient. This time series can be analyzed using classical Fourier spectral analysis resulting in spatiotemporal spectra for the gradient and curl flow (see Methods). Fig. 4c and d show changes in the spatiotemporal spectral power due to cortical lesioning (Fig. 4c) and following electrical stimulation in the acute phase after stroke (Fig. 4d) for both hemispheres. Most notably, changes in spectral power are different between the gradient and curl flow with the curl flow showing overall larger power increases across some temporal frequency bands particularly in the contralesional hemisphere. This suggests increases in feedback signaling following stroke and cortical stimulation as cortical feedback loops may give rise to circular communication patterns.

To demonstrate that the GDAR model can also be applied to field potentials from other recording modalities, we analyzed intracortical microelectrode array (Utha array) data from three sessions using either single-site or paired stimulation. For single site stimulation, we observe an increase in gamma GDAR flow power proximal to the stimulation site (Fig. 5b, left). For paired stimulations, a localized increase was only observed near the first stimulation site (Fig. 5b, middle and right), with no notable changes near the second site. We quantified this increase by computing the average flow magnitude over all edges connected to the stimulation site and adjusted it for changes in LFP power (Fig. 5c-e; also see Methods and Extended Fig. 4). Across all sessions, stimulation led to a sustained increase in resting-state flow near the first stimulation site above baseline levels. Notably, for paired stimulation sessions, this augmented flow persisted for at least 17 minutes following the final stimulation block.

#### The GDAR flow shows frequency specific correlations with reach velocity and exhibits directional tuning during a center-out reach task

Previous studies have shown that reach movements have strong neural correlates in M1 that can be detected from single neuron recordings as well as intracortical and surface field potentials^42–47^. Using µECoG recordings from M1 of a macaque monkey performing a center-out reach task (Fig. 6a), we show that the GDAR model can be used to study such neural correlates of behavior on the level of network FC dynamics. To do so, we leverage the graph spectral decomposition of the GDAR flow signal described above and in Methods to decompose the high-gamma GDAR flow into its gradient and curl flow spectrum (Extended Fig. 5a-e). An example of the time-varying flow spectra for a single reach trial along with the corresponding reach velocity is shown in Fig. 6b. Furthermore, using data from all directions and trials, Fig. 6c shows how the spectral power time series at each spatial frequency correlates with reach velocity. We found that an increase in reach velocity generally correlates with increases in gradient and curl flow power. Remarkably, the increase in gradient flow power is most pronounced only for the 15 lowest spatial frequencies. In contrast to higher frequencies, such low-frequency flow patterns are more coherent across the graph (see Extended Fig. 5b for examples), suggesting that coordinated activity across a larger cortical area facilitates reach movements.

**Fig. 6.**
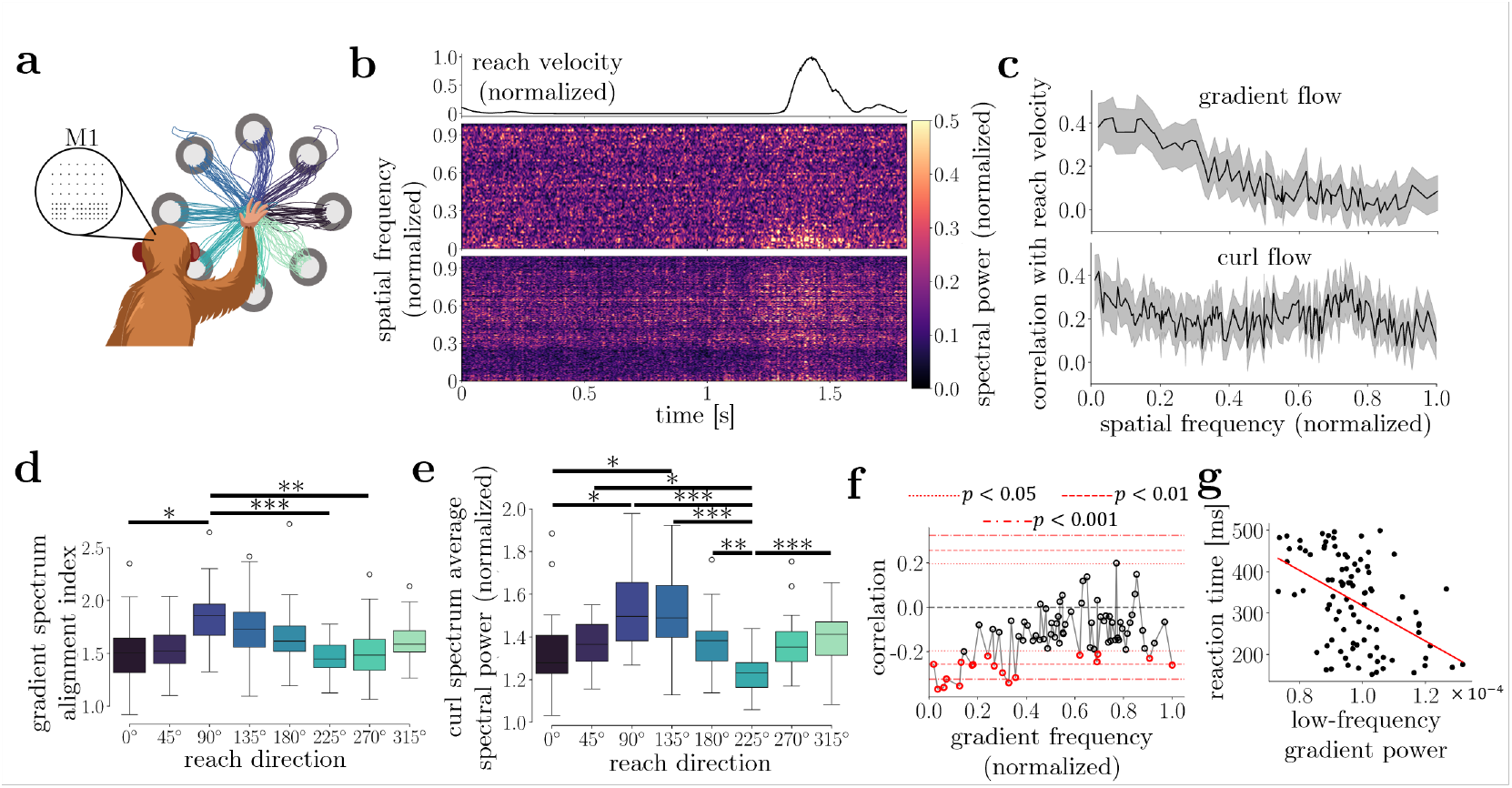
Applying the GDAR model to ECoG recordings during a center-out reach tasks. (a) A macaque monkey performed an eight directional reach tasks with 25 trials per direction while LFPs were recorded using a 96-channel µECoG array placed over the primary motor (M1) cortex. The GDAR flow was computed for each reach trial, bandpass filtered between 70-200 Hz and decomposed into its gradient and curl spectrum for each time bin (see Methods). (b) Gradient and curl spectrogram for a single reach trial. The black line shows the reach velocity. During the reach we observe an increase in curl flow power across all frequencies and gradient flow power for low spatial frequencies. (c) Correlation (median and interquartile range) between reach velocity and flow spectral power for all spatial frequencies pooled across all trials and directions. Low-frequency gradient flow components show the highest correlation with reach velocity suggestion more activity is coordinated across the M1 cortical network during reaching. Gradient alignment index, defined as the ratio of the 15 lowest to the 15 highest gradient flow spectral coefficients for all eight reach directions (shown are quartiles, 1.5 times the interquartile range, and outliers). The alignment index forms a cosine-like tuning curve with a preference for the 90° (up) and 135° (up-left) directions. (e) Same as in (d) but for the average curl flow power. (f) Correlation between average gradient spectral power during the last 100 ms before the go cue and the reaction time, defined as the time between the go cue and movement onset, for each gradient frequency. Especially low frequencies show a significant negative correlation with reaction time (g) Reaction time, as a function of the low gradient frequency spectral power during the last 100 ms before the go cue. The strong negative correlation suggests that more coordinated network activity leads to a faster reaction time (correlation coefficient: -0.483; p-value: 9.42·10^-7^).

To quantify the extent to which the gradient flow spectrum is dominated by low frequencies, we defined the gradient alignment index, which is computed as the ratio of the average power within the 15 lowest spatial frequencies to the average power within the 15 highest spatial frequencies (see Methods). The alignment index shows strong directional tuning with a preference for the 90° (up) and 135° (up-left) directions (Fig. 6d, and Extended Fig. 5g) and a similar cosine characteristic as reported in the literature for other recording modalities^42,43,46^. We also observe a similarly strong directional tuning characteristic for the average power of the curl spectrum (Fig. 6e and Extended Fig. 5h). In contrast, the directional tuning for the high-gamma envelope of the raw LFP signal averaged across all channels is significantly weaker (Extended Fig. 5i). This suggests that latent patterns of network activity extracted by the GDAR model rather than overall changes in signal power are better correlated with different behaviors.

Finally, we investigated if spectral network features derived from the GDAR flow are correlated to preparatory activity prior to movement onset. We found that both gradient and curl flow power computed during the last 100 ms prior to the go cue show negative correlations with the reaction time, which is defined as the time between go cue and initiation of movement, for most spatial frequencies (Fig. 6f and Extended Fig. 5j). This effect is most significant for low-frequency gradient modes (Fig. 6f and g) suggesting that a higher degree of neural coordination not only correlates with faster movements, but also indicates better movement preparation. We have shown that this effect cannot be explained by any premature movement activity (Extended Fig. 5k).

## Discussion

By drawing insights from computational neuroscience, statistical modeling, and signal processing over networks, we have introduced the GAR and GDAR model as members of a class of graphically constrained autoregressive models for estimating network-level dFC from field potential recordings. In contrast to classical VAR models that are a popular tool for electrophysiological FC analysis, our approach incorporates the spatial layout of the recording array as a structural prior, thereby significantly reducing the model complexity while mimicking cortical connectivity on a macroscopic scale. Furthermore, we demonstrate how our models give rise to a flow signal with the same temporal resolution as the original recordings, making them well suited for analyzing both transient and long-term FC patterns. Finally, network communication in the GDAR model is driven by gradients of neural activity—rather than activity itself as is the case for GAR and VAR models—resulting in a diffusion-like process. Using simulations and four electrophysiology datasets from macaque sensory motor cortex, we have demonstrated that the GDAR model outperforms competing approaches (GAR and standard VAR models) in estimating transient flow dynamics, provides complementary insights into resting state FC, and can uncover neural flow dynamics that correlate with behavior.

A drawback of standard VAR models is that they ignore spatial relations between the recording electrodes, which means that interactions between nearby sensors are treated equally to interactions between distant ones. However, especially for cortical recordings it is likely that signal propagation is dominated by short range connections due to the local connectivity structure of the cortex found in mice and macaque monkeys^23^. The idea of integrating spatial information in the form of structural priors into standard VAR models and other FC measures has recently been proposed in magnetic resonance imaging (MRI), electroencephalography (EEG), and magnetoencephalography (MEG) studies, where it has been shown to improve the estimation of FC networks^48–51^. However, to the best of our knowledge, it has not been applied to localized recording arrays that focus on network dynamics within one or two cortices. By adding structurally informed priors, the GAR and GDAR model have approximately five to ten times fewer parameters than the full VAR model for the electrode arrays used in our analysis resulting in less overfitting to idiosyncrasies in the data. The problem of overfitting is particularly evident in the poor generalization performance of the VAR model in simulations and for the reach dataset where model fitting relies on a limited number of observations (see Fig. 2f and Extended Fig. 2h).

Modeling dFC on top of a network also has the advantage that we can perform graph spectral analysis on the resulting dFC signal. This allows us to decompose complicated network-wide communication patterns into simpler and more interpretable modes representing feedforward (gradient) and feedback (curl) signal flow dynamics. Similar node-centric graph spectral decomposition techniques have recently been proposed to analyze neural data^52–57^, however these have exclusively focus on node signals rather than FC networks.

An important finding of our study is that the GDAR model outperforms the GAR model on simulated data even though the GAR model uses the same structural prior and is more expressive. Therefore, the better performance of the GDAR model must be attributed to the additional diffusion constraint incorporated into the model. Indeed, diffusive processes have previously been proposed as possible a mechanism for neural communication^29,30^. It has also been shown that diffusion processes can explain FC estimates^58^ and model the propagation of activity evoked by intracranial stimulation more accurately than alternative models of neural communication^59^. Furthermore, the Laplacian that drives the diffusion process in the GDAR model has been used in neural field models to simulate realistic large-scale brain dynamics^60,61^. Therefore, on a macroscopic level, the diffusion constraint may add a biophysically realistic mechanism to the model that aids in its overall performance. Nevertheless, we want to emphasize that the GDAR model is still a phenomenological model which is in contrast with dynamic causal modeling where the goal is to provide insights into biological mechanisms of neural communication^62–65^.

Using activity gradients as a driver of neural communication also has additional practical advantages for processing field potential signals. Such signals often suffer from spurious correlations due to volume conduction, signal artifacts that are shared across channels, and the common reference signal problem. These spurious correlations are known to cause erroneous connectivity estimates in many FC approaches such as coherence, phase locking value, or metrics based on standard VAR models^17^ and are commonly addressed by preprocessing field potentials using, common average referencing, current source density analysis (i.e., the second spatial derivative) or activity gradients (i.e., the first spatial derivative) instead of using the raw neural activity^17,66,67^. Since the GDAR model employs the second spatial derivative, a mitigation strategy against spurious correlations is naturally incorporated into the model making it more robust in a real-world setting.

Another key difference between existing dFC frameworks and our approach is that we propose a new mechanism for obtaining temporally resolved connectivity estimates. Existing approaches typically derive such dynamics through sliding windows^68–70^ or adaptive parameter estimation^51,71^. In contrast, we achieve this by combining static model parameter with the recorded neural activity resulting in a dynamic FC signal with maximal temporal resolution. This approach offers several advantages: it reduces the number of parameters that need to be estimated, and it enables the detection of short, transient communication events that may be smoothed out by sliding window approaches or are difficult to track using linear adaptive parameter estimation techniques. Using simulations, we also demonstrated that combining parameters with neural activity produces more accurate FC estimates than purely parameter-based techniques such as PDC and DTF (Fig. 2b and Extended Fig. 2d).

Our approach has several limitations. The assumption of a locally connected nearest neighbor graph as a structural prior inspired by the local connectivity structure of the cortex neglects the potential existence of any direct long-range propagation paths. Since it can be difficult to determine the best underlying network structure as precise structural information is often not available, we suggest that in the future the structural connectivity graph could be designed in a more data-driven way, for example, using sparsity and distance regularizers. Furthermore, we currently make no distinction between nodes corresponding to electrodes in the interior versus the boundary of the array. Especially the boundary nodes may exhibit a large exchange of information with regions outside the array, which is not captured by the model, but could be incorporated by adding additional input terms. Another promising avenue would be to explore how other proposed mechanisms of neural signaling, such as biased random walks or shortest path routing^29,30^, could be incorporated as constraints into data-driven dFC models. Furthermore, the GAR and GDAR model only capture linear interactions, but could be extended to model non-linear communication dynamics by introducing activation functions into the model.

Although we developed and tested our new framework using field potential recordings, we believe it can be extended to other neural recording modalities and applications. For instance, proposed models may be adapted for spiking data, either by modifying the autoregressive component to accommodate for discrete point processes—such as through generalized linear models^72^—or by first converting spikes into firing rates. The models should also be readily applicable to human ECoG and stereoelectroencephalography (sEEG) recordings, which share similar signal properties with the recordings analyzed here. Finally, GAR and GDAR models can be applied to brain-wide recoding modalities such as EEG, MEG or functional MRI (fMRI), which—combined with estimated structural connectivity networks—can enable the analysis of large-scale brain dynamics. In a preliminary investigation, we found that the GDAR model reliably estimates network-level dFC from resting-state fMRI data and is sensitive to age-related changes in neural flow^73^, highlighting its potential for broader applications in systems neuroscience and clinical research.

## Acknowledgements

We thank Daniel Silversmith and Joseph O’Doherty for their help with data collection and Philip Sabes for his laboratory in which some of the data was collected. We also thank Toni Haun, Sandi Thelen, and Christopher English for their help with animal surgeries and experimentation for the stroke experiment. This work was supported by the American Heart Association (FS, AY) the IEEE Signal Processing Society (FS), the National Institute of Health R01NS119593 (JB, AY) and R01MH125429 (FS, AY), the Washington Research Foundation (AY), the Big Data for Genomics and Neuroscience Training Grant NIH 5T32LM012419 (JB), the Center for Neurotechnology NSFERC 1028725 (JB), the Washington National Primate Research Center NIH P51OD010425, U42 OD011123 (AY), the Eunice Kennedy Shriver National Institute of Child Health and Human Development NIH K12HD073945 (AY), the National Institute of Neurological Disorders and Stroke of the National Institute of Health R01NS116464 (AY, JZ), the University of Washington Royalty Research Fund (AY, KK), the National Science Foundation Graduate Research Fellowship Program (KK), the Graduate Education for Minorities Fellowship (JB), and the Weill Neurohub (JZ).

## Methods

### Graph Autoregressive (GAR) and Graph Diffusion Autoregressive (GDAR) Model

#### Representation as constrained VAR model

Denoting the recorded neural activity at time *t* across all *N* recording channels as *s*[*t*]∈ ℝ ^*N*^, the classic vectorautoregressive (VAR) model can be written as

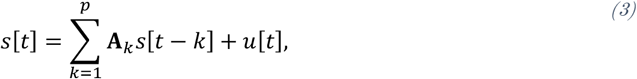

where *p* is the model order, **A***k* ∈ ℝ ^*N*×*N*^ contains the VAR model parameters, and *u*[*t*]∈ ℝ ^*N*^ is a white Gaussian noise term. The GAR model can be obtained by constraining the model parameters to be zero for connections not included in the structural connectivity graph 𝒢, that is (**A***k*)*i,j* = 0, if (*i, j*) ∉ 𝒢. The GDAR model is obtained by additionally enforcing a symmetry constraint on the matrix **A***k*, that is (**A***k*)*i,j* = (**A***k*)*j,i*, if (*i, j*) ∈ 𝒢. While this representation is most useful for estimating the model parameters from data, next we will provide an alternative and more intuitive interpretation for the GDAR model.

#### Derivation and Interpretation of the GDAR model as a network diffusion process

An alternative starting point for deriving the GDAR model is to describe the spatiotemporal dynamics of the neural activity *s*as a heat diffusion process

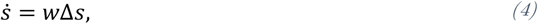

where temporal changes in activity 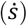 are driven by spatial activity gradients (Δ*s*, where Δ is the surface Laplacian) multiplied by the diffusion rate *w*. The right-hand side of (4) is equivalent to current source density (CSD) which is a common technique for analyzing neurophysiological recordings. The surface Laplacian Δ is equivalent to the second spatial derivative and thus describes local interactions within the brain network. In a discrete measurement setup with activity recorded from *N* recording channels, this can be encoded by constructing a locally connected graph from the locations of the electrodes within the recording array^74^. Thereby, each electrode corresponds to a node in the graph and edges connect neighboring electrodes such as illustrated in Fig. 1a. The resulting unweighted graph consisting of *N* vertices and *E* edges can be represented algebraically using the node-to-edge incident matrix **B** ∈ ℝ^*N*×*E*^, where the *e*^th^ column *b*^(*e*)^ corresponds to the *e*^th^ edge in the graph. Each edge is defined by a tail node *i* and head node *j* such that 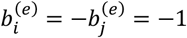 and all other entries 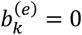. For each edge it is thereby arbitrary which incident node is defined as head and tail node. Using **B**, the continuous surface Laplacian Δ can be approximated using the negative of the graph Laplacian operator **B**^*T*^. Furthermore, the first temporal derivative 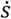 can be approximated by the first temporal difference *s*[*t*] − *s*[*t* − 1]. Thus, (4) can be approximated by

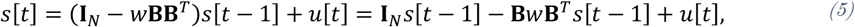

Where, **I**_*N*_ is the *N* × *N* identity matrix and *u*[*t*] is a white noise term. Previously, it has been shown that the matrices **B**^*T*^ and **B** can be interpreted as discrete approximations of the gradient and divergence operators, respectively^26^. Thus, the term **B***w***B**^*T*^*s*[*t* − 1] has a clear physical meaning in the context of LFP recordings:

1. **B**^*T*^*s*[*t* − 1] = ∇*s*[*t* − 1]: computes the voltage gradient for each node in the graph.
2. *w*∇*s* [*t* − 1] = *f* [*t*]: In analogy to resistive circuits and CSD analysis, *w* can be interpreted as a conductivity such that conductivity times voltage gradient yields a current flow *f*[*t*].
3. **B***f*[*t*]: For each node, the net flow, i.e., the sum of all inflows minus the sum of all outflows, is computed. This is equivalent to computing the current sources and sinks in CSD analysis.

Equation (5) effectively expresses CSD analysis as a first order vector autoregressive (VAR) model. However, the model in Eq. (5) has limited expressivity as the only learnable parameter is the conductivity *w*. Thus, to improve its expressivity, we can 1) add parameterized node dynamics, 2) assume a spatially varying conductivity, and 3) extend the model order from one to *p*. The resulting graph diffusion autoregressive (GDAR) model is given by

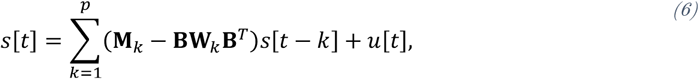

where **M**_*k*_ = diag(***m***_*k*_) ∈ ℝ^*N*×*N*^ and **W**_*k*_ = diag(***w***_*k*_) ∈ ℝ^*E*×*E*^ are diagonal matrices containing the node and edge parameters ***m***_*k*_ ∈ ℝ^*N*^ and ***w***_*k*_ ∈ ℝ^*E*^ of the *k*^th^ lag, respectively. The term **BW**_*k*_**B**^*T*^ can also be regarded as a weighted graph Laplacian matrix. The GDAR flow is defined as

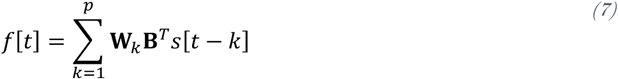

It can be shown that the GDAR model in (6) can be rewritten to match the standard notation of a VAR model in Eq. (3) by setting (**A**_*k*_)_*i, j*_ = (**A**_*k*_)_*j, i*_ = (**W**_*k*_)_*l,l*_ if *l* corresponds to the edge between node *i* and *j* and (**M**_*k*_)_*i,i*_ = (**A**_*k*_)_*i, i*_ + ∑_*j*∈*𝒩*(*i*)_(**A**_*k*_)_*i,j*_, where *𝒩*(*i*) are the set of neighbors of node *i*.

### Model fitting

Using the VAR representation in Eq. (3), the model parameters **M**_*k*_ and **W**_*k*_ can be estimated using least squares regression following the procedure described by Lütkepohl^75^. Given *T* + *p* snapshots of neural activity by an *N*-channel recording array, where *T* is the number of samples used for model fitting, the predicted neural activity can be collected in the data matrix **Y** = *s*[*p* + 1], … , *s*[*p* + *T*] ∈ ℝ^*N*×*T*^ and its vectorized version *γ* = vec(**YY**). The regressors can be expressed as **S** = [*S*_1_, … , *S*_*T*_] ∈ ℝ^*Np*×*T*^, where *S*_*t*_ = [*s*[*t* + *p* − 1]^*T*^, … , *s*[*t*]^*T*^]^*T*^ ∈ ℝ^*Np*×1^. The coefficients **A**_*k*_ can be expressed as **A** = [**A**_1_, … , **A**_*p*_] ∈ ℝ^*N*×*Np*^ and *α* = vec(**A**). As shown in the previous section, **A**_*k*_ is spare for the GAR and sparse and symmetric for the GDAR model. Therefore, there exist a matrix **R** such that 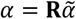 and 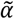 only contains the non-zero entries of **A** (GAR) or the non-zero upper triangle entries of **A** (GDAR). Now (3) can be written as

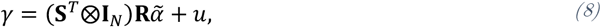

where ⨂ is the Kronecker product. Furthermore, we assume that *u* is white noise with covariance matrix **Σ**_*u*_. Eq. (8) can be solved in close form by minimizing 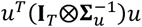 , where **I**_*T*_ is the *T* × *T* identity matrix, to obtain the optimal parameters 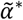 :

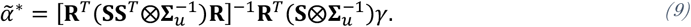

Eq. (9) is the solution to the generalized least squares (GLS) estimator, which is generally different from the ordinary least squares (OLS) estimator due to the sparsity and symmetry contraints^75^. However, it requires knowledge of the noise covariance matrix **Σ**_*u*_, which is unknown in practice. Therefore, we first estimate the **Σ**_*u*_ by solving the OLS estimator *u*^*T*^*u* to compute 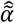 as

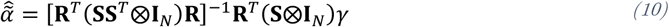

and denote 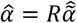. The corresponding coefficient matrix is **Â** with 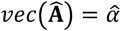. Then we estimate **Σ**_*u*_ as

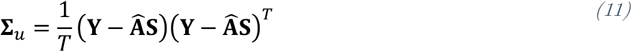

It is also noted that for the GDAR model, Eq. (6) can be directly cast as a least squares minimization problem. However, we found that it is more efficient to compute the optimal parameters according to (8).

### Power spectrum of GAR and GDAR flow

If the models are applied to resting state neural activity, the GAR and GDAR flow may exhibit a similar oscillatory behavior as the neural activity. Therefore, it may be reasonable to compute its power spectrum to study frequency specific dFC patterns. Using Eq. (7) and recognizing that it expresses the flow as the convolution between the model parameters **W**_*k*_ and the activity *s*[*t*] (activity gradients **B**^*T*^*s*[*t*] for GDAR), the flow power spectrum between nodes *i* and *j* is given by

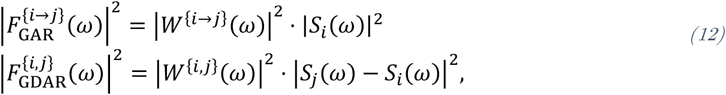

where *W*^{*i* , *j*}^(*ω*), *S*_*i*_ (*ω*), and *S*_*j*_(*ω*) are the Fourier transforms of the parameters, as well as the neural activity of the two channels, and *ω* is the frequency variable. An interesting case occurs for the GDAR model when the spectra of both channels have the same magnitude for a given frequency. Assuming |*S*_*i*_ (*ω*)| = *S*_*j*_(*ω*) = 1, Eq. (12) can be simplified to

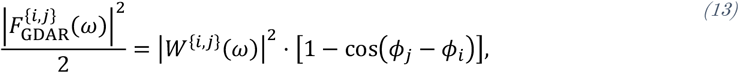

where *Φ*_*i*_ and *Φ*_*j*_ are the phase of the *S*_*i*_ (*ω*), and *S*_*j*_(*ω*), respectively. That is, in this case the GDAR flow dynamics are driven only by phase differences between connected nodes. In general, however, GDAR flow dynamics will be determined by differences in magnitude and phase modulated by *W*^{*i* , *j*}^(*ω*), which was estimated with the objective of predicting future neural activity.

### Decomposition into gradient and curl flow spectra

Similar to the classical Fourier transform for time series, where a signal can be decomposed into a series of oscillatory components of increasing frequency, a flow signal can be decomposed into a set of spatial components (flow signals) with increasing spatial frequency. Furthermore, a flow signal can be decomposed into gradient (non-circular) components, which have non-zero divergence (sum of in-flow minus out-flow) for some or all nodes of the graph, and curl components, which have zero divergence for all graph nodes. This can be achieved via the Hodge-decomposition that defines two orthogonal sets of spatial basis functions (defined on the edge domain) for a given graph^25–27^. A directional graph flow signal, such as the GDAR flow at each timestep *t*, can then be projected onto these sets of basis functions to obtain the gradient and curl flow spectrum. Note that this decomposition assumes a unidirectional flow *f* ^{*i* , *j*}^ for each edge in the graph, where the sign of *f* ^{*i* , *j*}^ together with the edge reference direction defined in **B** determines the flow direction. For bidirectional flow signals *f* ^{*i* →*j*}^ ≠ *f* ^{*j*→*i*}^ such as for the GAR or VAR models, we first need to obtain a unidirectional flow, which we generally define as *f* ^{*i* , *j*}^ = *f* ^{*i* →*j*}^ − *f* ^{*j*→*i*}^

To obtain the gradient basis, we first compute the eigenvectors 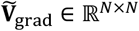 of the graph Laplacian **B**^*T*^. The orthonormal gradient flow basis **V**_grad_ ∈ ℝ^*E*⨯*N*^ is then obtained by

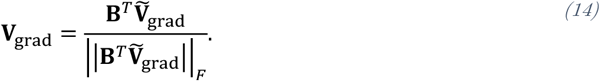

The eigenvalues *λ*_grad_ associated with each eigenvector define a natural ordering of the eigenvectors in terms of spatial frequency. Specifically, if we compute the divergence of the eigenvectors **V**_grad_, we find that eigenvectors corresponding to small eigenvalues have small divergence, whereas eigenvectors associated with large eigenvectors have large divergence. Small-divergence eigenvectors correspond to flow signals that are *smooth* (or low-frequency) across the graph, that is flow signals where the direction of flow is largely preserved or only slowly changes within a local neighborhood (see Extended Fig. 1a for an example). High-divergence eigenvectors on the other hand correspond to flow patterns that rapidly change direction within a local neighborhood and can therefore be considered as *non-smooth* or being high-frequency. We can now obtain a gradient flow spectrum for each flow snapshot by projecting *f* [*t*] onto **V**_grad_:

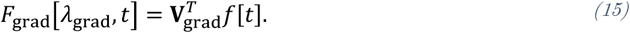

To obtain the curl basis, we first must define a set of triangles in the graph, which can be obtained, for example, by finding all possible triangles in the graph. Mathematically, the triangle set is captured by the edge-to-triangle incident matrix **B**_tri_ ∈ ℝ^*E*⨯*T*^ , where *T* is the number of triangles and where the *t*^th^ column 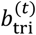 corresponds to the *t*^th^ triangle in the graph. Each triangle is defined by three edges *e*_*i*_ , *e*_*j*_, and *e*_*k*_ and an arbitrarily chosen reference direction. If the edge direction across *e*_*i*_ (as defined in **B**) aligns with that reference direction 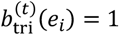. Otherwise 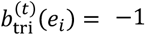 (the same logic applies to *e*_*j*_ , and *e*_*k*_). For edges not involved in the triangle we have 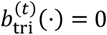. To compute the curl basis, we then follow the same procedure as above. That is, we first compute the eigenvectors 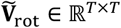 of the Laplacian 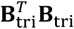 and then project 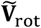 onto **B**_tri_ and normalize:

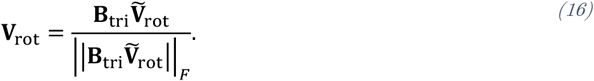

Similar to the gradient flow, the eigenvalues *λ*_rot_ corresponding to the eigenvectors **V**_rot_ can be used to define an ordering in terms of spatial frequency. Specifically, eigenvectors with small eigenvalues correspond to global curl flows (akin to global currents) across the graph that maintain or only slowly change orientation between local neighborhoods. On the other hand, eigenvectors with small eigenvalues exhibit localized curl flows (akin to local eddy currents) that rapidly change orientation across local neighborhoods (see Extended Fig. 1b for an example). Finally, we can obtain a curl flow spectrum for each flow snapshot by projecting *f* [*t*] onto **V**_rot_:

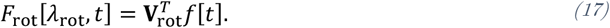

Finally, temporal changes in the time-varying gradient and curl flow spectra can be analyzed using classical Fourier-based spectral analysis techniques, resulting in spatiotemporal spectra 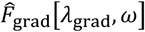, and 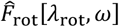, where *ω* denotes the temporal frequency.

### Wilson-Cowan Simulations

#### Simulating neural activity

We simulated neural activity using various networks of Wilson-Cowan oscillators^76,77^ shown in Fig. 2 and Extended Fig. 2. Each node consists of an excitatory and inhibitory subpopulation whose dynamics are governed by the following differential equations:

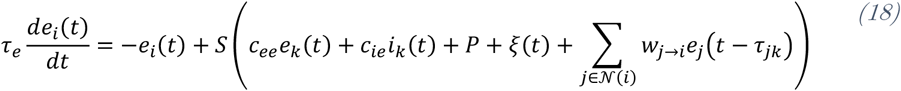

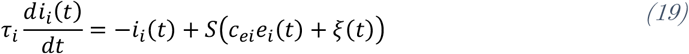

where *S* is the sigmoid function:

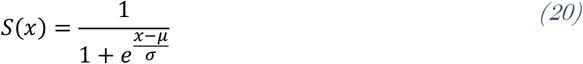

The description of the parameters and their values are listed in Table. The values are based on previous work by Abeysuriya et al.^78^ and Deco et al.^79^ and result in a power spectrum with a pronounced beta oscillation around 18 Hz and a 1⁄ω slope for higher frequencies. Coupling between excitatory populations of neighboring nodes is determined by the parameter *wj*→*i*where each edge in the graph has two coupling parameters (*wj*→*i*and *wi*→*j*) resulting in bidirectional coupling. For the 16-node random graphs, we simulated 10 independent trials per graph, resulting in a total of 100 trials for 10 graphs, where for each trial the values of the edge weights *wj*→*i*are randomly sampled from a uniform distribution (see Table 1 for range of *wj*→*i*). For the 7-node graph, we simulated 100 independent trials respectively. The ranges of *wj*→*i*were chosen such that neural activity whose power spectrum resembles realistic local field potential signals was generated by the network. We integrated the system with a time step of 0.1 ms seconds using a 4th order Runge-Kutta scheme for 20 seconds and discarded the first 5 seconds to eliminate transient effects of the simulation. The resultant 15 seconds of excitatory activity *e*[*t*]was then downsampled to 1 kHz using an 8th order Chebyshev type I anti-aliasing filter and denoted as the simulated neural activity. Power spectral density (PSD) estimates of the simulated activity for the 16-node random graphs averaged over all trials, graphs, and edges are shown in Extended Fig. 2b.

**Table 1.** Simulation parameters of Wilson-Cowan model adapted from Abeysuriya et al.^78^ and Deco et al.^79^.

| Parameter | Description | Value |
| --- | --- | --- |
| $\tau_e$ | Excitatory time constant | 0.002 |
| $\tau_i$ | Inhibitory time constant | 0.004 |
| $c_{ee}$ | Local excitatory to excitatory coupling | 3.5 |
| $c_{ie}$ | Local inhibitory to excitatory coupling | -2.5 |
| $c_{ei}$ | Local excitatory to inhibitory coupling | 3.75 |
| $P$ | Constant excitatory input | 0.31 |
| $\mu$ | Firing response threshold | 1 |
| $\sigma$ | Firing threshold variability | 0.25 |
| $\xi$ | Random noise input | $\mathcal{N}(0, 0.05)$ |
| $w_{i \rightarrow j}$ | Excitatory to excitatory connectivity (16 node random graphs) | 0.05, ..., 0.3 |
| $w_{i \rightarrow j}$ | Excitatory to excitatory connectivity (7 node graph) | 0.05, ..., 0.55 |

#### Simulating ground truth neural flow

We simulated the ground truth flow by calculating the moment-to-moment influence that each excitatory node exerts on its neighbors. To do so, we first executed each integration step with the full set of parameters to obtain *e*[*t*]. Then, for each excitatory coupling parameter *wj*→*i*, we repeated the integration step with *wj*→*i*= 0 to obtain 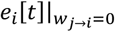, which denotes the activity at node *i*in the absence of an influence from node *j* at time *t*. The flow from node *j* to node *i*was then defined as 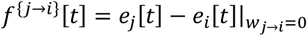. This second step is repeated for all excitatory coupling parameters. The full two-step procedure is repeated for each integration step. The resulting bidirectional ground truth flow *f*gt,b[*t*]was downsampled using the same anti-aliasing filter as used for the simulated neural activity. As our GDAR model only produced a unidirectional flow at each point in time, we define the unidirectional ground truth flow *f* [*t*]between node *i*and *j* as 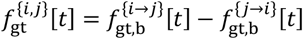. It is noted that while *f*gt[*t*]is unidirectional at each time point, the flow direction across each edge can change over time and is determined by the sign of *f*gt[*t*]. PSD estimates of the unidirectional ground truth flow for the 16-node random graphs averaged over all trials, graphs, and edges are shown in Extended Fig. 2c.

#### VAR, GAR and GDAR flow

For each trial, we used the last 15 seconds of simulated neural activity to estimate the parameters of VAR, GAR and GDAR models for varying model orders as described above. The graph used for fitting the model is equal to the graph used for the simulations. The estimated model parameters were then used to transform the simulated neural activity into an estimated flow signal according to Eq. (2). For the VAR and GAR models, the unidirectional flow is defined as 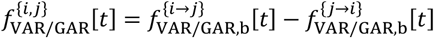. Since the VAR model assumes a fully connected graph, we generally obtain non-zero flow signals across connections that are not part of the network. For fair comparison with the ground truth flow, we only extract the VAR flow for edges that exist in the ground truth network. For the GAR models, we also compute a bidirectional flow *f*GAR,b[*t*]and compare it to the ground truth bidirectional flow *f*gt,b[*t*]. However, we found that this does not result in higher correlation coefficients than the unidirectional flow (see Extended Fig. 2e).

#### Partial directed coherence (PDC) and directed transfer function (DTF)

GAR, GDAR, and VAR models (order *p* = 24) were also used to compute PDC^22^ and DTF^21^ from the model parameters, which were then correlated with the ground truth flow spectra. Traditional PDC and DTF estimates (Fig. 2b) were computed from the VAR model parameters for each directed edge pair *j* → *i*using the following equations:

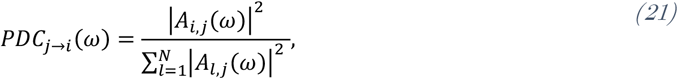

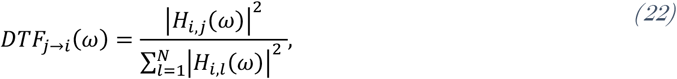

where *Ai,j*(ω) is the Fourier transform of (*Ak*)*i,j* (note that here *k* is the time variable), and *Hi,j* (ω) is the (*i, j*) entry of the Fourier transform of the inverse of *Ak*. Equivalent metrics can be obtained for the GAR and GDAR models by computing the sum in the denominator only over the nodes connected to *j* and *i*respectively (Extended Fig. 2d). For the GDAR model, the estimated PDC and DTF spectra are compared to the flow spectra obtained from the unidirectional ground truth flow (see above). For GAR and VAR models, bidirectional PDC and DTF estimates are obtained and compared to the bidirectional ground truth flow spectra.

#### Comparing ground truth and estimated neural flow

The ground truth and estimated flow signals are first z-scored independently for each trial and model and then compared using various metrics. First, power spectral density (PSD) estimates of the flow across each edge was estimated using Welch’s method^80^ with a Hann window of size 256 samples and 50% overlap and PSDs of the ground truth and estimated flow signals were correlated using Pearson correlation coefficients and compared between models using Wilcoxon rank-sum tests (Fig. 2b and d). Furthermore, correlation coefficients between raw time series of ground truth and estimated neural flow were computed and pooled across all edges and trials (Extended Fig. 2f). Next, we computed the error between magnitude and phase spectrum of ground truth and estimated flow for each graph edge and trial (Fig. 2c). Magnitude errors are defined as the absolute difference (in dB) between ground truth and estimated flow PSD. To compare the phase error, the 15s of neural flow obtained for each trial were first divided into 58 non-overlapping segments of 256 samples and then the discrete Fourier transform for each segment was computed. Afterwards, the phase difference between ground truth and estimated phase was computed and mapped into the range from 0 to π for each segment before being averaged over all 58 segments. Finally, we compared the dynamics of the estimated flow with the dynamics of the ground truth flow using dynamical similarity analysis (DSA) in Fig. 2e^33^. To do so, Hankel dynamic mode decomposition (DMD) models are first independently fitted to the high-dimensional ground truth and estimated flow signal and the resultant DMD matrices **A**_est_ and **A**_gt_ are compared using a modified version of Procrustes analysis. To fit the Hankel-DMD models we used 15 delay time steps to construct the Hankel matrices and full rank regression. Optimization during the Procrustes analysis used 1000 iterations at a learning rate of 10^−2^.

#### Generalization performance

According to Eq. (6), the GDAR model can predict neural activity at the current time step using the past *p* samples. To assess the generalization performance of the model, we computed the normalized root mean square error (RMSE) between the observed neural activity *s*[*t*]and predicted neural activity Ŝ [*t*]as follows:

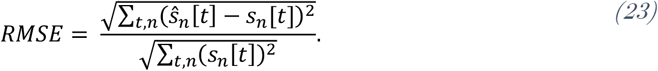

The summation is performed over all time points *t* within a segment, as well as over all nodes *n*. To compute the train RMSEs, the predictions Ŝ [*t*]are computed for the same time points that were used for model fitting. For the test RMSEs, the models are applied to data immediately following the training set. For the simulations in Fig. 2f, RMSEs were evaluated using 100 independent trials from the 7-node graph shown in Extended Fig. 2g. We used 10^th^ order models and the length of train and test segments were set equal. The generalization gap is defined as the difference between test and train RSME after averaging over all edges and trials.

To compute the generalization gap for the in-vivo experiments (Extended Fig. 2h), the models were fitted to 10 s training segments (optogenetic stimulation, stroke, and Utah array) and evaluated on the subsequent 10 s segment. For the reach dataset, the models are tested on the subsequent reach trial in the same direction. test RMSEs for the optogenetic stimulation, stroke, and Utah array datasets described above, the prediction RMSEs are computed for the 10 s segment that immediately follows the segment used for model fitting. For the center-out reach experiment, the same data as described above was used. For the optogenetic stimulation, Utah array, and reach experiment the same data as described below were used. For the stroke experiment in addition to the data described below, an additional 30 min of baseline recording at the end of the experiment was used.

### Optogenetic Stimulation Experiment

One adult male rhesus macaque (monkey G: 8 years old, 17.5 kg) was used in this experiment. All procedures were performed under the approval of the University of California, San Francisco Institutional Animal Care and Use Committee and were compliant with the Guide for the Care and Use of Laboratory Animals.

#### Neural stimulation and recording interface

In this study, we used a subset of neural data recorded by a large-scale optogenetic neural interface^11^ that has previously been utilized to study changes in network FC due to cortical stimulation^34,35^. The interface was composed of several key components: a semi-transparent micro-electrode array, a semi-transparent artificial dura, a titanium implant, and a laser system for delivering optical stimulation. First, neurons in the primary sensorimotor cortex were rendered light-sensitive through a viral-mediated expression of the C1V1 opsin. To do so, 200 µL of the viral cocktail AAV5-CamKIIa-C1V1(E122T/E162T)-TS-eYFP-WPRE-hGH (2.5 × 10^12^ virus molecules/mL; Penn Vector Core, University of Pennsylvania, PA, USA, Addgene number: 35499) was administered across four sites into the primary somatosensory (S1) and primary motor (M1) cortices of the left hemisphere using convection-enhanced delivery^11,34,81^. Next, the chronic neural interface was surgically implanted by performing a 25mm craniotomy over the primary sensorimotor cortex, replacing the dura mater beneath the craniotomy with a chronic transparent artificial dura housed in a titanium chamber. During each experimental session, the artificial dura was removed and a custom 96 channel micro-electrocorticography array consisting of platinum-gold-platinum electrodes and traces encapsuled in Parylene-C^12^ was placed on the cortical surface. Optical stimulation was performed by two 488 nm lasers (PhoxX 488-60, Omicron-Laserage, Germany) connected to a fiber optic cable (core/cladding diameter: 62.5/125 μm, Fiber Systems, TX, USA) and positioned above the array such that the tip of the fiber-optic cable touched the array. Neural data in the form of local field potentials was recorded by the micro-ECoG array at a sampling frequency of 24 kHz using a Tucker-Davis Technologies system (FL, USA). It was verified that evoked neural responses were due to optogenetic activation and not other effects such as photoelectric artifacts or heating^11,12,35^.

#### Stimulation protocol

For the stimulation-evoked analysis, we used data from three experimental sessions all performed on the same day. The only difference between the sessions was the location of stimulation, which is depicted in Fig. 3b. As the micro-ECoG array was not removed between sessions its location on the cortex remains unchanged. Each experimental session consists of 5 stimulation blocks during which two lasers alternatingly and repeatedly stimulate. Each stimulation block lasts approximately 7 min and is intermittent by shorter resting state blocks during which no stimulation is performed. The stimulation pulse width for both lasers was 5 ms with a delay of 10 ms between stimulation by lasers 1 and 2. This paired stimulation is repeated at a frequency of 7 Hz (143 ms) resulting in a total of approximately 2970 pulse pairs for each stimulation block. All stimulation parameters (except for stimulation locations) are identical for the three sessions analyzed in Fig. 3.

#### Signal preprocessing

First, bad channels were identified as 1) electrodes with high impedance and 2) channels with a low signal-to-noise ratio, and excluded from the analysis^35^. The location of the remaining 67 good channels was used to construct a sparse and locally connected graph, where each electrode corresponds to a node in the graph and each node is connected approximately to it 8 nearest neighbors (see Fig. 3c top). The raw time series data was downsampled to 1017.25 Hz using a low-pass Chebyshev anti-aliasing filter and the mean activity within each channel was subtracted from the respective time series.

#### GDAR model fitting

The preprocessed LPFs during each stimulation block were divided into segments of 10004 samples (approximately 10 s) with 4 samples overlap between segments and a 5^th^ order GDAR, GAR, and VAR models were fitted to each segment as described above. The estimated model parameters were used to transform the neural activity into a flow signal according to equation (7). The overlap between segments was chosen such that a continuous flow signal was obtained from the segmented LFPs without relying on zero padding. A model order of 5 was chosen for this application due to the short (10 ms) delay between stimulation by lasers 1 and 2. For larger model orders, the flow evoked by the second laser would increasingly be influenced by the neural activity evoked by the first laser resulting in a mixing of the neural responses to both stimulation pulses. Flow dynamics akin to the plots in Fig. 3d for a model order *p* = 10 are shown in Extended Fig. 3c and d.

#### Visualizing flow dynamics

To visualize the flow dynamics evoked by paired cortical stimulation, we must project the high dimensional flow signals *f*[*t*]∈ ℝ^*E*^, where *E* is the number of edges in the graph, for the GAR and GDAR flow onto a lower dimensional subspace. To so so, we first pooled the first 25 flow snapshots from the onset of stimulation by the first laser from all sessions, blocks, and pulse pairs in a single data matrix **F** ∈ ℝ^*E*×*M*^, where *R* ≈ 3 ⋅ 5 ⋅ 2970 ⋅ 25 (3 sessions, 5 blocks per session, approximately 2970 pulse pairs per block, 25 flow snapsots per pulse pair). Afterwards we performed principle component analysis (PCA) and projected **F** onto its first two principal components (PCs) to obtain 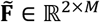. Fig. 3d and e show the PCA reduced GDAR, and GAR flow dynamics where each trace illustrates a 25 snapshot long flow trajectory from a single pulse pair. For better visualization only 250 individual trajectories per stimulation block selected at random are plotted. Fig. 3f shows the same dynamics but instead using the VAR flow. Since the number of edges for the VAR model is very large, computing the PCs of the associated matrix *R* was not feasible. Therefore we first averaged the flow snapshots over 20 consequitive trials before computing the PCs (resulting in *M* ≈ 3 ⋅ 5 ⋅ 148 ⋅ 25, where 148 is the number of averaged trials). For comparison, we performed the same trial averaing for the GDAR flow and recomputed the flow trajectories (Extended Fig. 3e). The averaging does not seem to have a negative effect on the discriminability of the trajectories between different sessions.

#### Modeling increased delay across sulcus

The GDAR model can easily be augmented to model variable delay across different edges. For example, it is reasonable to assume that signals that travel across the sulcus between M1 and S1 experience larger delays than signals traveling within each cortex. Larger delays in the GDAR model across an edge between node *i*and *j* can be incorporated by constraining edge coefficients 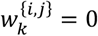 for small delays (i.e., *k* = 1, 2, …), which can be achieved by augmenting the matrix **R** in equation (8). We have used this approach to model larger delays across the sulcus by setting 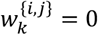 for edges that connect nodes in M1 to nodes in S1 for *k* = 1, 2, 3. That is, the minimum delay across each sulcus edge is constrained to be 4 (see Extended Fig. 3f and Supplementary Video 2 for corresponding GDAR flow dynamics).

### Changes in resting-state FC

To demonstrate the GDAR model’s ability to uncover changes in FC during resting state, we analyze data from a stroke and cortical stimulation experiment using two 32-channel ECoG arrays, and an intracortical stimulation experiment using a 96-channel microelectrode array (Utah array).

#### Stroke and cortical stimulation experimental procedure

One adult macaque (*Macaca nemestrina*, 14.6 kg, 7 years, male) was used in this study. All procedures were performed under the approval of the University of Washington Institutional Animal Care and Use Committee and were compliant with the Guide for the Care and Use of Laboratory Animals. The experimental procedure was previously described elsewhere^13,41,82^. The animal was first anesthetized with isoflurane and a craniotomy with 25mm diameter was performed in each hemisphere over the sensorimotor cortex. A focal ischemic lesion in the left hemisphere was created by photo-activation of a previously injected light-sensitive dye (Rose Bengal). Following illumination, the dye causes platelet aggregation, thrombi formation, and interruption of local blood flow, leading to local neural cell death near the illuminated area. The location and extent of the lesion were estimated through post-mortem histological analysis of coronal slices and is illustrated as a black patch in Fig. 5. Electrical activity was recorded before (30 min), during (60 min), and after lesion induction (60 min) simultaneously in the ipsi- and contralesional hemisphere using two 32-channel ECoG arrays (Fig. 5c)^83,84^. Approximately 60 min after the end of lesioning, repeated electrical stimulation was performed 8 mm away from the lesion center. 1 kHz stimulation charge-balanced pulses (60 µA, 450 µs pulse width, 50 µs interphase interval) were given in 5 Hz bursts (5 pulses per burst) consecutively for 10 minutes, where each stimulation block was followed by 2 min of baseline recording. The experiments included a total of six 10 min stimulation blocks. We used the 30 min of neural recordings before lesion induction, 60 min of neural recording after lesion induction but before stimulation (pre stim), as well as the 2 min blocks of baseline recording in between the stimulation blocks (post stim) in both hemispheres. In total we used 4 blocks of post stimulation recordings as the recordings in the other blocks were corrupted. All data were digitally notch-filtered at 60, 180, and 300 Hz using the function scipy.signal.iirnotch with -3 dB bandwidth of 3 Hz.

#### Intracortical stimulation experimental procedure (Utah array)

One adult rhesus macaque (*Macaca mulatta*, 12 kg, 11 years, male) was used in this study. All procedures were performed under the approval of the University of California, San Francisco Institutional Animal Care and Use Committee and were compliant with the Guide for the Care and Use of Laboratory Animals. The experimental procedure was previously described by Bloch et al.^85^. A 96-channel Utah array was implanted in S1 and LFPs were recorded at a sampling frequency of 24 kHz before being downsampled to a frequency of 1017 Hz (8^th^ order Chebychev anti-aliasing filter). The dataset consists of resting state recordings intermitted by five 10 min stimulation blocks that contain repeated single site or paired electrical stimulation. For the single site stimulation session, stimulation is performed in in the form of five pulses (1 kHz burst frequency) that are repeated every 200 ms. The paired stimulation sessions use the same stimulation patterns for each stimulation site. For session *paired-stim 1*, electrode B stimulated 100 ms after electrode A. For session *paired-stim 2*, the delay between stimulation sites A and B is chosen uniformly at random between -100 ms and 100 ms for each paired stimulation trial.

#### Signal preprocessing

The preprocessing for both datasets was performed akin to the optogenetic stimulation experiment. The location of the ECoG channels was used to construct a sparse and locally connected graph, where each node (electrode) is connected approximately to its 8 nearest neighbors. When necessary, the raw time series data were downsampled to 1 kHz using low-pass Chebyshev anti-aliasing filter and the mean was removed from each channel. Additionally, artifacts—defined as signal values that deviate by ten or more standard deviations from the mean simultaneously for all channels—were removed by linearly interpolating between the sample immediately before and after the artifact for the stroke dataset.

#### Model fitting and postprocessing

The preprocessed LFPs for all datasets during each block were divided into 10,009 sample long segments (approximately 10 s) with 9 samples overlap between segments and 10^th^ order GDAR and and VAR models were fitted to each segment as described above. For the GDAR model, the estimated model parameters were used to transform the neural activity into a flow signal according to equation (2), where each segment contains 10000 samples. To assess FC changes in different frequency bands, we then computed the GDAR flow power spectral density (PSD) using Welch’s method^80^ (Hann window of size 1000 samples with 50% overlap) for each segment, and stored the average flow PSD within the gamma band (30 − 70 Hz). Finally, we computed the change in average flow PSD due to stroke and stimulation. Specifically, if we denote 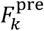 and 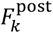 as the average gamma flow PSD before and after stroke/stimulation of the *k*^th^ segment, the relative change in GDAR flow magnitude Δ_stim_ is given by

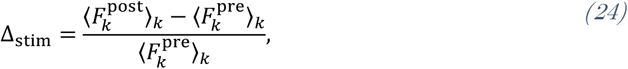

where ⟨⋅⟩_*k*_ denotes the average over all segments. We assessed the statistical significance of Δ_stim_ for each edge by forming sample distributions for the pre and post condition from all segments and compared the distributions using a two-sample Kolmogorov-Smirnov test. If the distributions for a given edge differ with a significance level of *p* ≤ 0.01, the edge is plotted in the graph. For partial directed coherence (PDC) and directed transfer function (DTF), the same procedure is followed with PDC and DTF spectra being computed over each 10-second segment according to equation (21) and (22) using the VAR model parameters.

Spatiotemporal spectra 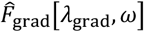 and 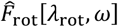 in Fig. 4c and d were first calculated for each segment of the pre and post condition as described above using Welch’s method^80^ (Hann window of size 1000 samples with 50% overlap) for the temporal Fourier analysis. Afterwards spectral power was calculated by taking the magnitude square of 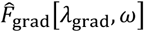 ,and 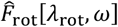 and averaged over all segments.

For the Utah array data, changes in gamma GDAR flow power due to stimulation (Fig. 5b) were computed using all data before stimulation (pre stim) as well as the five 2-minute resting state blocks following the stimulation blocks (post stim) for each session. To compute the temporal evolution of the normalized and averaged gamma GDAR flow power (Fig. 4c-e), the GDAR flow power in the gamma band was first averaged over all edges connected to the stimulation node. Then GDAR flow and LFP power were z-scored using the mean and standard deviation from the pre stim period for each session independently. We then computed the best linear fit between the z-scored LFP and average GDAR flow power 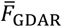 using all segments (pre and post stim; Extended Fig. 4)

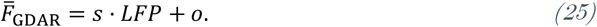

The goal is to test whether the GDAR flow power changes beyond what can be linearly explained by changes in LFP power. Hence, we subtract the linear regression line from the average GDAR flow power

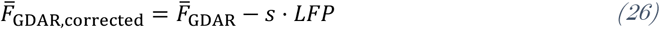

and plotted the result in Fig. 4c-e.

### Center-out Reach Task

One adult male rhesus macaque (7 years old, 16.5 kg) was used in this study. All procedures were performed under the approval of the University of California, San Francisco Institutional Animal Care and Use Committee and were compliant with the Guide for the Care and Use of Laboratory Animals. Surgical procedure, neural interface, and signal preprocessing are the same as described in Optogenetic Stimulation Experiment. However, for the center-out reach task, the ECoG was placed fully over the primary motor (M1) cortex. Channels with persistent distortions were identified and excluded from the analysis resulting in 77 good channels used to construct a sparsely connected nearest neighbor graph as described previously. The animal performed a total of 200 successful reach trials, 25 for each of the eight directions (see Fig. 6a). Each individual reach trial is divided into start, instructed delay, and reach phase. During the start phase, the monkey places its hand on the center of the screen. After that the instructed delay phase begins where first the target direction is presented before a randomly selected delay period terminated by a go-tone is introduced. The reach phase starts once the go-tone appears and ends when the monkey touches the target. The finger position of the monkey was tracked throughout the experiment using and electromagnetic position sensor (Polhemus Liberty, Colchester, VT) at 240 Hz^86^.

#### GDAR model fitting and post-processing

To ensure accurate model fitting, recorded LFPs from all three phases were used to estimate the parameters of the GDAR model. The model order was set to *p* = 5 to ensure enough independent samples for each parameter. After the model parameters have been estimated, the GDAR flow is computed according to Eq. (7) and filtered into the high-gamma band using a 3^rd^ order Butterworth filter with cutoff frequencies of 70 and 200 Hz. The high-gamma GDAR flow signal *f*[*t*]is then decomposed into its gradient and curl flow spectrum *F*_*grad*_λ_*grad*_, *t* and *F*_rot_[λ_rot_, *t*]according to Eq.(14)-(17). To obtain the flow power spectrogram in Fig. 6b, we compute the magnitude square of *F*_*grad*_λ_*grad*_, *t* and *F*_rot_[λ_rot_, *t*]. Flow power spectra as well as reach velocities are temporally smoothed using a 51 sample 3^rd^ order Savitzky-Golay filter. To account for the time delay between motor commands observable in M1 and actual movement onset^87^, we calculated the median correlation across all gradient frequencies between *F*_*grad*_λ_*grad*_, *t* − *d* and the reach velocity for varying delays *d* (Extended Fig. 5f). We found a maximum correlation for a delay of 104 ms, which we corrected for in all subsequent analysis.

To quantify the extent to which the gradient flow spectrum is dominated by low frequencies during reaching, we defined the alignment index as

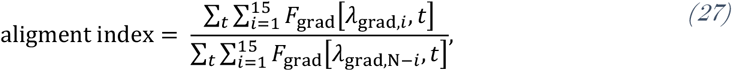

where *N* is the total number of gradient frequencies. The temporal averaging is performed over all time points where the reach velocity is above a threshold of 0.1 for the tuning curve analysis (Fig. 6d and e). For the curl flow spectrum, we do not observe spectral changes during reaching that are strongly dependent on the spatial frequency. Therefore, we simply use the average over all spatial frequencies in Fig. 6e.

The reaction time is defined as the time between the go cue and when the finger displacement reaches 5mm (approximately 10% of the full path length) from the resting position during the delay period. This reaction time is then correlated with the average GDAR flow power for each spatial frequency over the last 100 ms prior to the go-tone (Fig. 6f and Extended Fig. 5j). Furthermore, we have excluded trials with a reaction time smaller than 150 ms and larger than 500 ms resulting in a total of 93 valid trials. The correlation in Fig. 6g uses the average GDAR flow power over the five lowest gradient frequencies.

## Data availability

Data will be made available upon reasonable request from the authors.

## Code Availability

Source code for the GDAR model will be made available prior to publication.

## Supplementary Notes

### Modeling signal propagation delay across sulcus for optogenetic stimulation experiment

The GDAR model can also be adapted to model longer signal propagation paths between specific nodes in the network. For example, this may be the case for connections across the sulcus that separate M1 and S1. To model this, we have constrained the GDAR model to enforce a minimum propagation delay of 4 ms for all edges that connect nodes in M1 with nodes in S1 (see Methods). The evolution of the GDAR flow for Session 1 averaged over all trials is shown in Supplementary Video 2 and is similar to the dynamics observed in Fig. 3d and Supplementary Video 1, but with noticeably less flow across the sulcus. The PC reduced flow dynamics across all sessions are shown in Extended Fig. 3f and are almost identical to the dynamics shown in Fig. 3e, suggesting that adding signal propagation constraints to the model does not negatively impact the sensitivity of the GDAR flow signal to the stimulation parameters.

## Extended data figures

**Extended Fig. 1.**
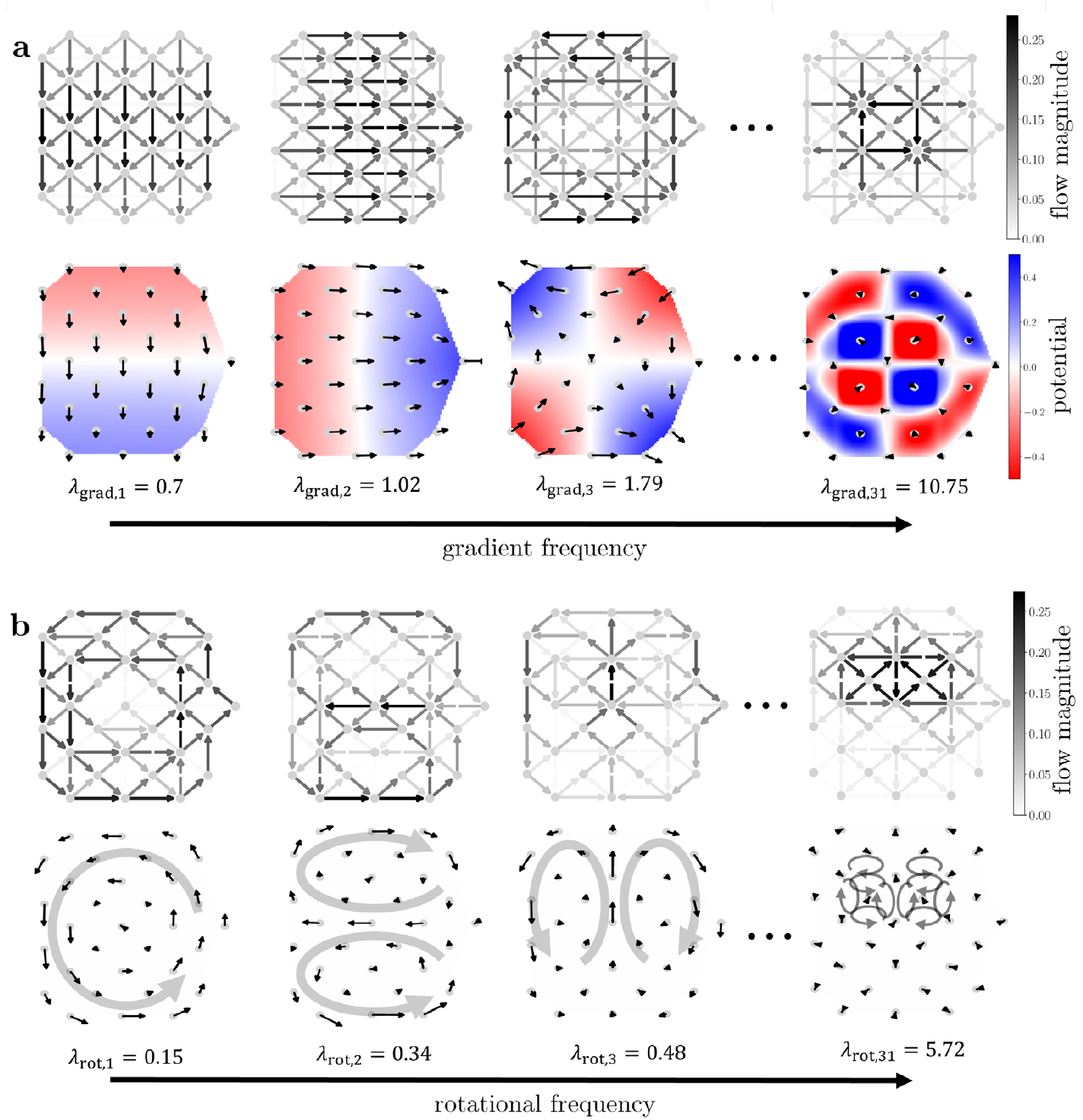
Basis functions for flow spectral decomposition of the structural connectivity graph used in the stroke experiment. (a) First three and last gradient basis function V_grad_ and their corresponding eigenvalue describing non-circular flow patterns. The top row shows the basis function as a flow signal on top of the structural connectivity graph. To better visualize the spatial frequency characteristics, we compute the flow fields (net-flow magnitude and direction at each node) and potential fields (divergence of flow at each node) for each basis function (bottom row). The eigenvalues can be interpreted as spatial frequencies with basis functions corresponding to larger eigenvalues exhibiting more localized and chaotic flow patterns. (b) Same as (a) but for the curl basis functions V_rot_ describing circular flow patterns. The gray arrows in the flow fields are manually drawn for better visual identification of the dominant flow patterns. Gradient and curl flow spectra are computed by projecting a flow signal f onto the corresponding basis functions (top rows of (a) and (b)).

**Extended Fig. 2.**
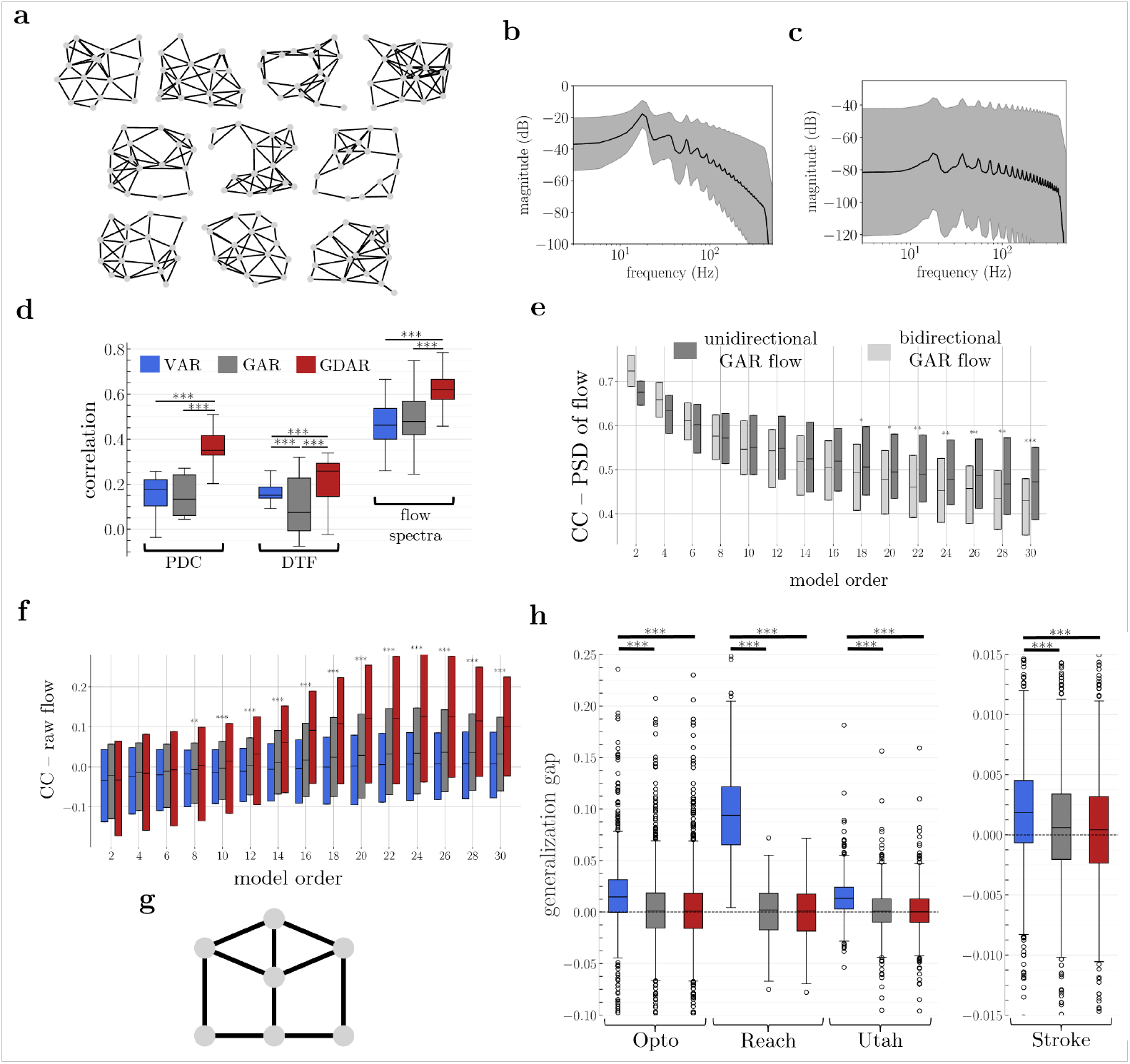
(a) 10 randomly connected 16-node networks used for conducting the simulations in Fig. 2. Each graph was used to generate 10 independent simulation trials. (b), (c) Power spectral densities of simulated field potentials and ground truth flow, respectively. The black line shows the average across all edges and trials. The gray shaded area indicated one half of the standard deviation. The simulation parameters produce a strong oscillation around 18Hz. The steep drop-off above 400 Hz can be attributed to the 8^th^ order Chebyshev filter that was used for downsampling the data to a sampling frequency of 1 kHz. (d) Distribution of Pearson correlation coefficients (CC) (medians, upper and lower quartiles, interquartile range) between ground truth flow spectrum and partial directed coherence (PDC) and directed transfer function (DTF) computed using VAR (traditional), GAR and GDAR model. Correlation with flow spectrum (same as in Fig. 2b) is shown for reference as well. (e) CC between power spectral density (PSD) of unidirectional ground truth and GAR flow, as well as bidirectional ground truth and GAR flow for various model orders. For model orders p ≥ 18, the unidirectional GAR flow yields significantly higher median CCs than the bidirectional GAR flow. (f) CC between time-domain ground truth and estimated neural flow for varying model orders. Markers indicate whether the GDAR model significantly outperforms the GAR model (Wilcoxon ranked-sum test, p ≤ 0.05) The GDAR model significantly outperforms all other models for orders p ≥ 8. (g) 7-node graph used for assessing the generalization performance of the VAR, GAR, an GDAR model on simulated data. (h) Generalization gap of all models for four electrophysiology datasets. GAR and GDAR model almost perfectly generalize to unseen data. On the other hand, the VAR model always shows some degree of overfitting (Wilcoxon rank-sum test, p ≤ 0.001).

**Extended Fig. 3.**
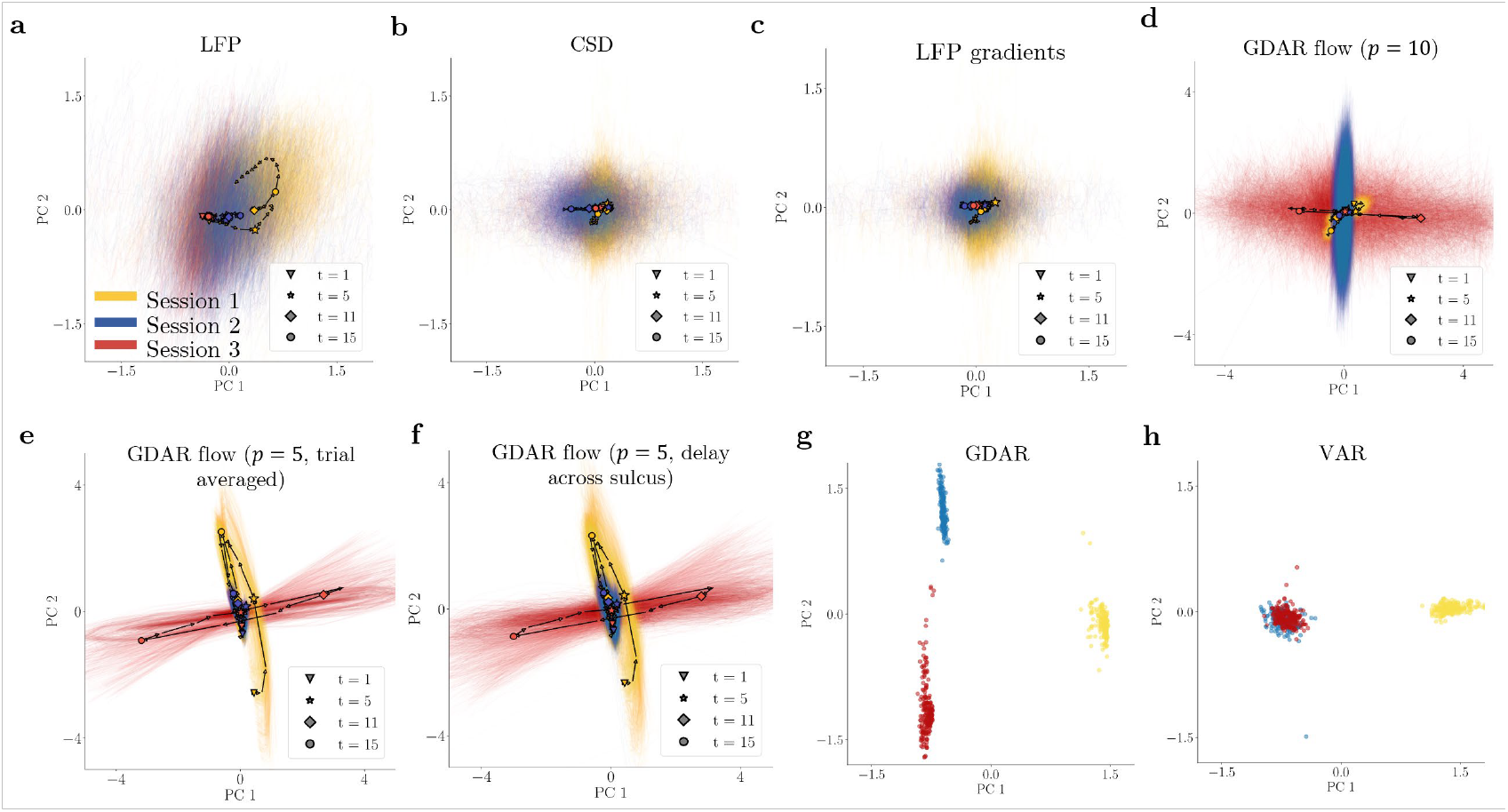
(a)-(f) Stimulation evoked dynamics using different signals and modeling approaches similar to the plots in Fig. 3e. Neither LFP (a), classical CSD (b), i.e., the second spatial derivative computed via the graph Laplacian, nor LFP gradients (c) show significant temporal dynamics that are distinct between the sessions. (d) Using a 10^th^ order GDAR model results in reduced separability between sessions when compared to a 5^th^ order model shown in Fig. 3e. This is likely a result of the time scale of the paired stimulation (the onset of the second laser pulse occurs 5 ms after the offset of the first pulse) causing a mixing of the effects of the two lasers when fitting the models. (e) Averaging the flow signal over 20 consecutive trials before computing the low dimensional embedding does not have a negative effect on the dynamics. (f) PC reduced GDAR flow dynamics when constraining edges crossing the sulcus between M1 and S1 to exhibit a minimum signal propagation delay of 4ms. The dynamics are almost identical to the dynamics from the unconstrained GDAR model (Fig. 3e), suggesting that the stimulation induced communication dynamics uncovered by our model are robust to such constraints. (g), (h) The parameters of the 5^th^ order GDAR and VAR model for all segments, blocks, and sessions were stacked into a single matrix and projected onto its first two PCs. Each dot represents the PC reduced parameters of a single 10 s segment used for model fitting. The parameters from the three different sessions are well separated in this low dimensional subspace for the GDAR model, but not for the VAR model where Session 2 and 3 are not separable.

**Extended Fig. 4.**
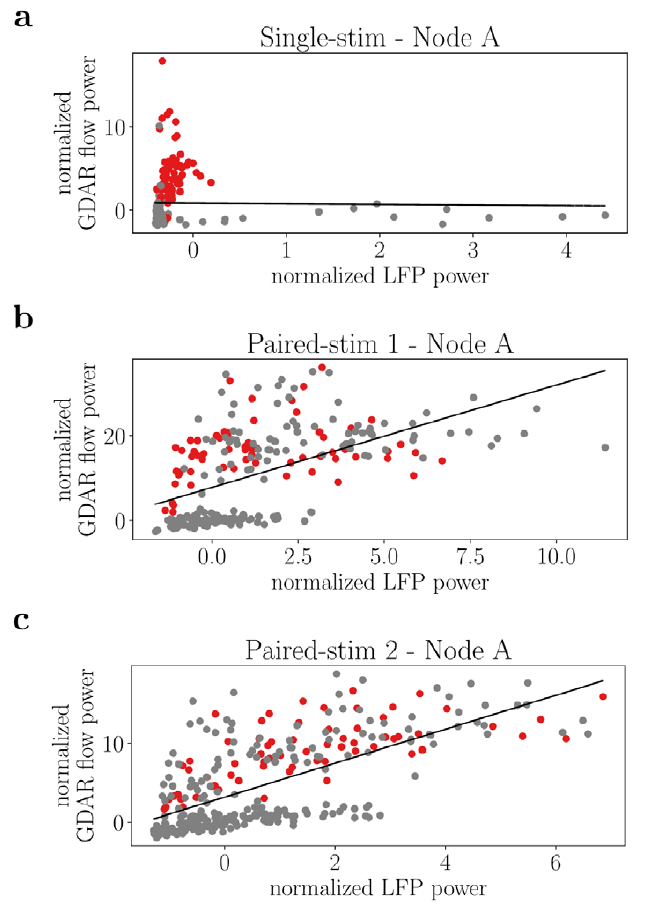
Relation between average gamma GDAR flow and LFP power for the stimulation electrodes in the Utah array dataset. The best linear regression lines are also shown (Single-stim: slope = −0.0692, p = 0.827; Paired-stim 1:slope = 2.41, p = 1.05e − 19; Paired-stim 2: slope = 2.15, p = 2.1e − 43). The two paired stim sessions show a strong linear relation between LFP and average GDAR flow power. Nevertheless, the GDAR flow power increase due to stimulation is larger than what can be explained by linear changes in LFP power.

**Extended Fig. 5.**
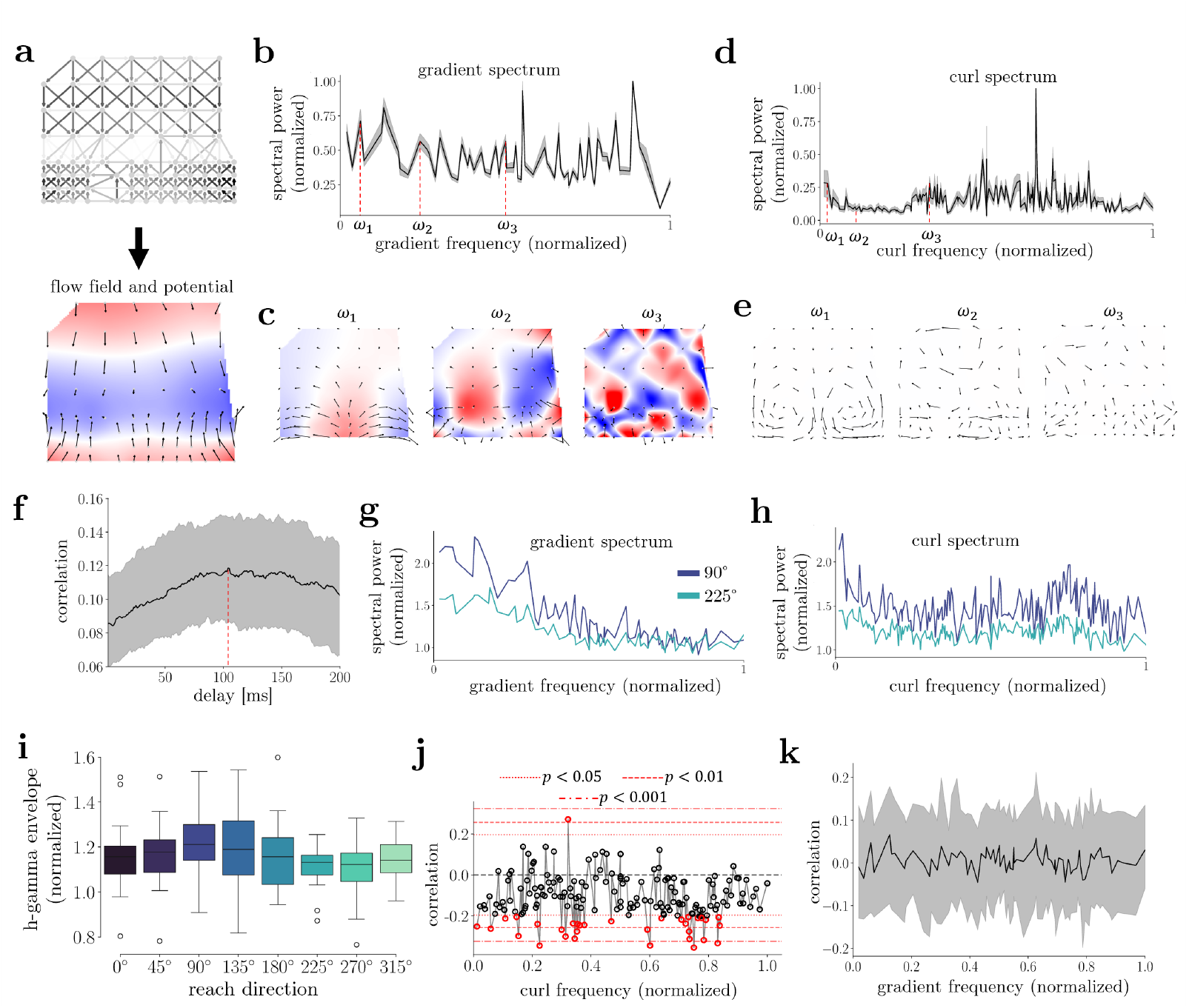
Extended plots for reach data shown in Fig. 6. (a) Example gradient flow component as well as flow and potential field corresponding to the 6^th^ lowest spatial frequency. (b) Average spectrum of the gradient flow component (median and interquartile range) across all time points, trials and directions. Note that the spectrum largely resembles white noise with a few stronger spectral components at the low, mid, and high frequencies. (c) Flow and potential fields for three spectral components marked in (b). As the frequency increases, the flow becomes more disorganized with an increasing number of local sources and sinks and the overall divergence (measure of spatial frequency) increases. (d) and Same as (b) and (c), but for curl flow. The average curl flow spectrum is primarily marked by an increase in power for some high-frequency components. (f) Correlation (median and interquartile range) between average gradient flow power (averaged over all spatial frequencies) and reach velocity for different delays between the neural signal and the recorded velocity. The maximum correlation occurs at 104 ms, which we assume to be transmission delay between motor commands in the brain and observable movements. (g) and (h) Average gradient and curl power spectra for the 90° (up) and 225° (bottom-left) directions. The 90° direction shows a substantially larger increase in the low frequency gradient flow power, as well as curl flow power across almost all frequencies than the 225° direction, highlighting the directional tuning of the GDAR flow. (i) Directional tuning curves (quartiles, 1.5 times interquartile range, and outliers) for envelope of high gamma (70 − 200 Hz) filtered local field potential signal averaged over all recording electrodes. The same trend as for the GDAR flow alignment index (Fig. 6d) and average curl flow power (Fig. 6e) can be observed, however differences between directions are not significant. (j) Correlation between average curl spectral power and reaction time for each curl frequency. (k) Correlation (median and interquartile range) between average gradient flow power during the last 100 ms prior to the go-cue and reach velocity 104 ms later. The median correlation of zero suggests that there is no residual movement occurring during that period that could explain the correlation between the alignment index and the reaction time in Fig. 6g.

## Supplementary Material

**Supplementary Table 1.** p-values (one-sided Wilcoxon rank-sum tests) comparing GDAR and GAR model for differences in Pearson correlation coefficients (PCC) of the flow power spectral density (PSD; Fig. 2d), dynamical similarity analysis (DSA) dissimilarity scores (Fig. 2de), and PCC of the raw flow (Extended Fig. 2f). We used one-sided Wilcoxon rank-sum tests with the alternative hypothesis that the GDAR model outperforms the GAR model. Furthermore, p-values for comparing PSD correlations between the bidirectional and unidirectional GAR flow (Extended Fig. 2e) are shown. We used one-sided Wilcoxon rank-sum tests with the alternative hypothesis that the bidirectional outperforms the unidirectional GAR flow.

| Model order | PCC – PSD of flow | DSA dissimilarity | PCC – raw flow | GAR bidirectional vs. GAR unidirectional |
| --- | --- | --- | --- | --- |
| 2 | 0.59 | 0.373 | 0.982 | 0.999999977 |
| 4 | $3.8 \cdot 10^{-4}$ | 0.157 | 0.51 | 0.9998 |
| 6 | $1.39 \cdot 10^{-5}$ | 0.397 | 0.0694 | 0.933 |
| 8 | $2.6 \cdot 10^{-6}$ | 0.728 | 0.00148 | 0.613 |
| 10 | $2.69 \cdot 10^{-6}$ | 0.916 | $8.36 \cdot 10^{-7}$ | 0.307 |
| 12 | $1.66 \cdot 10^{-6}$ | 0.99 | $2.15 \cdot 10^{-13}$ | 0.103 |
| 14 | $8.53 \cdot 10^{-8}$ | 0.99939 | $5.82 \cdot 10^{-33}$ | 0.105 |
| 16 | $4.66 \cdot 10^{-10}$ | 0.999995 | $2.11 \cdot 10^{-57}$ | 0.0775 |
| 18 | $1.39 \cdot 10^{-11}$ | 0.9999988 | $5.76 \cdot 10^{-60}$ | 0.0345 |
| 20 | $8.95 \cdot 10^{-15}$ | 0.9999967 | $3.02 \cdot 10^{-69}$ | 0.0279 |
| 22 | $1.28 \cdot 10^{-14}$ | 0.987 | $1.35 \cdot 10^{-61}$ | 0.00511 |
| 24 | $1.13 \cdot 10^{-15}$ | 0.806 | $6.82 \cdot 10^{-66}$ | 0.00556 |
| 26 | $1.42 \cdot 10^{-12}$ | 0.536 | $4.65 \cdot 10^{-71}$ | 0.00263 |
| 28 | $1.34 \cdot 10^{-11}$ | 0.147 | $3.38 \cdot 10^{-67}$ | 0.00223 |
| 30 | $1.92 \cdot 10^{-9}$ | 0.134 | $4.04 \cdot 10^{-64}$ | 0.000544 |

**Supplementary Table 2.**
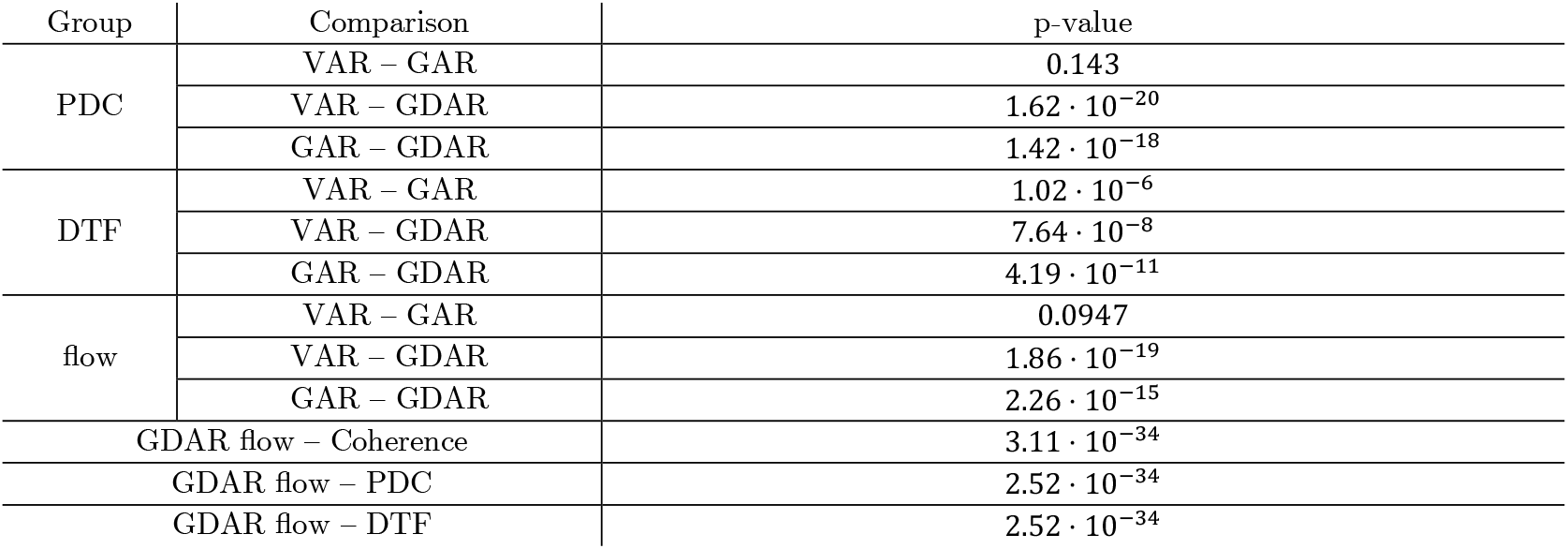
p-values (two-sided Wilcoxon rank-sum tests) for comparisons in Fig. 2b and Extended Fig. 2e. all comparisons in Fig. 2d. Shown are the within group comparisons partial directed coherence (PDC), directed transfer function (DTF), and flow power (Extended Fig. 2e), as well as the comparison between GDAR flow power and coherence, VAR-based PDC, and VAR-based DTF (Fig. 2b; p-values for those comparisons not marked in figure.)

| Group | Comparison | p-value |
| --- | --- | --- |
| PDC | VAR – GAR | 0.143 |
| | VAR – GDAR | $1.62 \cdot 10^{-20}$ |
| | GAR – GDAR | $1.42 \cdot 10^{-18}$ |
| DTF | VAR – GAR | $1.02 \cdot 10^{-6}$ |
| | VAR – GDAR | $7.64 \cdot 10^{-8}$ |
| | GAR – GDAR | $4.19 \cdot 10^{-11}$ |
| flow | VAR – GAR | 0.0947 |
| | VAR – GDAR | $1.86 \cdot 10^{-19}$ |
| | GAR – GDAR | $2.26 \cdot 10^{-15}$ |
| GDAR flow – Coherence | | $3.11 \cdot 10^{-34}$ |
| GDAR flow – PDC | | $2.52 \cdot 10^{-34}$ |
| GDAR flow – DTF | | $2.52 \cdot 10^{-34}$ |

**Supplementary Table 3.** p-values (one-sided Wilcoxon rank-sum test) for tuning curves in Fig. 6d and e. Only significant comparisons are shown. All p-values are corrected for multiple comparisons via the Bonferroni method. 0° and 90° correspond to the right and top direction, respectively.

| Gradient spectrum alignment index (Fig. 6e) | Curl spectrum average spectral power (Fig. 6f) |
| --- | --- |
| 90° - 0°: 0.0165 | 0° - 90°: 0.0165 |
| 90° - 225°: $4.42 \cdot 10^{-5}$ | 0° - 135°: 0.0165 |
| 90° - 270°: 0.00111 | 45° - 225°: 0.0165 |
| | 90° - 225°: $7.04 \cdot 10^{-6}$ |
| | 135° - 225°: $4.42 \cdot 10^{-5}$ |
|  | 180° - 225°: 0.00455 |
| | 315° - 225°: $4.42 \cdot 10^{-5}$ |

**Supplementary Table 4.** p-values (two-sided Wilcoxon rank-sum test) for differences in generalization gap between GAR, GDAR and VAR model for four electrophysiological datasets in Extended Fig. 2h.

|  | Opto | Reach | Rest Utah | Stroke |
| --- | --- | --- | --- | --- |
| VAR - GAR | $3.51 \cdot 10^{-15}$ | $2.65 \cdot 10^{-60}$ | $4.72 \cdot 10^{-39}$ | $1.19 \cdot 10^{-6}$ |
| GDAR - GAR | 0.768 | 0.598 | 0.735 | 0.329 |
| GDAR - VAR | $8.27 \cdot 10^{-16}$ | $1.76 \cdot 10^{-60}$ | $4.05 \cdot 10^{-40}$ | $1.07 \cdot 10^{-8}$ |

